# Leveraging auxiliary data from arbitrary distributions to boost GWAS discovery with Flexible cFDR

**DOI:** 10.1101/2020.12.04.411710

**Authors:** Anna Hutchinson, Guillermo Reales, Thomas Willis, Chris Wallace

## Abstract

Genome-wide association studies (GWAS) have identified thousands of genetic variants that are associated with complex traits. However, a stringent significance threshold is required to identify robust genetic associations. Leveraging relevant auxiliary covariates has the potential to boost statistical power to exceed the significance threshold. Particularly, abundant pleiotropy and the non-random distribution of SNPs across various functional categories suggests that leveraging GWAS test statistics from related traits and/or functional genomic data may boost GWAS discovery. While type 1 error rate control has become standard in GWAS, control of the false discovery rate can be a more powerful approach. The conditional false discovery rate (cFDR) extends the standard FDR framework by conditioning on auxiliary data to call significant associations, but current implementations are restricted to auxiliary data satisfying specific parametric distributions, typically GWAS *p*-values for related traits. We relax these distributional assumptions, enabling an extension of the cFDR framework that supports auxiliary covariates from arbitrary continuous distributions (“Flexible cFDR”). Our method can be applied iteratively, thereby supporting multi-dimensional covariate data. Through simulations we show that Flexible cFDR increases sensitivity whilst controlling FDR after one or several iterations. We further demonstrate its practical potential through application to an asthma GWAS, leveraging various functional genomic data to find additional genetic associations for asthma, which we validate in the larger, independent, UK Biobank data resource.

**Author summary:** Genome-wide association studies (GWAS) detect regions of the human genome that are associated with various traits, including complex diseases, but the power to detect these genomic regions is currently limited by sample size. The conditional false discovery rate (cFDR) provides a tool to leverage one GWAS study to improve power in another. The motivation is that if two traits have some genetic correlation, then our interpretation of a low but not significant *p*-value for the trait of interest will differ depending on whether that SNP shows strong or absent evidence of association with the related trait. Here, we describe an extension to the cFDR framework, called “Flexible cFDR”, that controls the FDR and supports auxiliary data from arbitrary distributions, surpassing current implementations of cFDR which are restricted to leveraging GWAS *p*-values from related traits. In practice, our method can be used to iteratively leverage various types of functional genomic data with GWAS data to increase power for GWAS discovery. We describe the use of Flexible cFDR to supplement data from a GWAS of asthma with auxiliary data from functional genomic experiments. We identify associations novel to the original GWAS and validate these discoveries with reference to a larger, more highly-powered GWAS of asthma.

## Introduction

Genome-wide association studies (GWAS) identify risk loci for a phenotype by assaying single nucleotide polymorphisms (SNPs) in large participant cohorts and marginally testing for associations between each SNP and the phenotype of interest. Conducting univariate tests for each SNP in parallel presents a huge multiple testing problem for which a stringent *p*-value threshold is required to confidently call significant associations.

The statistical power to detect associations can be increased by leveraging relevant auxiliary covariates. For example, pervasive pleiotropy throughout the genome ^1^ suggests that leveraging GWAS test statistics for related traits may be beneficial, whilst the non-random distribution of trait-associated SNPs across various functional categories ^2^ suggests that incorporating functional genomic data may also be useful. In fact, the expansive range of relevant auxiliary covariates has accumulated in a wealth of covariate-informed multiple testing methods which leverage auxiliary covariates (e.g. SNP-level data) with test statistics for variables (e.g. GWAS *p*-values for SNPs) to increase statistical power. These methods have been extensively researched both in the statistical literature^3,4,5,6,7,8,9,10,11,12,13^ and specifically in the context of GWAS^14,15,16,17,18,19,20,21,22^ with a consistent aim of minimizing type-2 errors (or equivalently increasing statistical power) whilst controlling some appropriate type-1 error rate, such as the false discovery rate (FDR).

The conventional multiple testing correction procedures control some error measure by assuming that each hypothesis is equally likely *a priori* to be true or false. For example, the Bejamini-Hochberg (BH) procedure ^23^ provides nearly optimal control of the FDR under the condition that almost all null hypotheses are true ^10^. The simplest extension to incorporate covariates is independent filtering ^9^ whereby test statistics are first censored based on the value of the covariate and a multiple testing method (for example, the BH procedure) is then applied on the remaining subset of test statistics. Alternatively, the test statistics can be grouped based on covariate values and the BH procedure can be applied separately within each group, as in stratified FDR ^4^. However, these simplistic approaches do not make full use of the information contained within the covariate values and require subjective covariate thresholding which may lead to data dredging bias ^6^.

Insinuating a more unified approach of incorporating covariates, the conventional multiple testing correction procedures can incorporate weights for each variable, such that the raw test statistics are no longer considered exchangeable ^24,25^. However, it is non-trivial to convert covariate values to weights satisfying certain constraints required for type-1 error rate control (typically non-negative weights that average 1^19^), and consequently the covariate values are typically still only used in an initial stratification step. For example, the grouped Benjamini-Hochberg (GBH) procedure ^5^ groups test statistics based on covariate values and derives group-specific weights that are proportional to the estimated proportion of true nulls in each group, whilst the independent hypothesis weighting (IHW) method calculates optimal group-specific weights which maximise the number of discoveries, whilst controlling the FDR ^6^. In the GWAS setting, the FINDOR method (which was shown to be generally less powerful, but superior in terms of false positive findings, to stratified FDR, GBH and IHW methods) ^15^ groups SNPs based on how well they tag functional categories that are enriched for heritability ^26,27^ and derives group-specific weights for use in a weighted Bonferroni procedure ^3^ that are proportional to the ratio of the estimated proportion of alternative to null SNPs in each group ^5,28^. The FINDOR methodology is similar to that of the GBH procedure, but includes an additional step whereby the weights are normalised to average 1. This normalisation step is significant because Roeder et al. ^19^ demonstrated that using a data-dependent weighting scheme with weights that average to 1 preserves control of type-1 error with high probability if the number of weights learned is significantly less than number of hypothesis test performed ^15^. Whilst grouping-based approaches are satisfactory when they capture all information provided by the covariate, as is possible in the case of categorical covariates, this assumption is restrictive in the case of continuous covariates or more complex multi-dimensional covariate spaces. Moreover, subjective thresholding and coarse binning is generally required, meaning that the entire dynamic range of the auxiliary data is often not fully exploited (for example, applying FINDOR to Biobank style data with the recommended 100 bins results in bins containing approximately 100K SNPs within which covariate values will vary).

Other notable methods in the GWAS literature include those which require subjective thresholding, such as GenoWAP ^14^ which thresholds on the covariate value to define “functional” SNPs, or those which require additional knowledge, such as the distribution of the true effect size^19,20,21^. Related methods in the statistical literature include those which estimate the proportion of true null hypotheses conditional on observed covariates and use these as plug-in estimates for the FDR ^29^, such as Boca and Leek’s FDR regression ^30^, those utilising the local FDR (defined as the posterior probability that a variable is null given the observed test statistic ^31^)^32,7,33^ and those that focus power on more promising hypotheses ^34,11,35,10^. Specifically, the AdaPT method ^10^ adaptively estimates a Bayes optimal *p*-value rejection threshold, but this requires a SNP filtering stage in the GWAS context due to computational demand ^36^. A comprehensive overview of these statistical methods for covariate-informed multiple testing is provided by Ignatiadis and Huber^37^.

The conditional FDR (cFDR) approach developed by Andreassen and colleagues ^38,39,40^ is a natural extension to the FDR in the presence of auxiliary covariates. This intuitive approach mitigates many of the aforementioned shortcomings: it does not bin variables and thus makes full use of the dynamic range of covariate values, it does not include any subjective thresholding, and does not require the definition of a normalised weighting scheme. However, it was designed for a very specific setting, that is to increase GWAS discovery (in the “principal trait”) by leveraging GWAS test statistics from a genetically related (“conditional”) trait.

We were interested to see whether a more general form of cFDR could address the same covariate-informed multiple testing problems as the range of methods listed above. Here, we describe “Flexible cFDR”, a new cFDR framework ^41^ that enjoys all of the benefits of the conventional approach but now supports continuous auxiliary data from arbitrary distributions, thus enabling broader applicability. Specifically, our computationally efficient approach extends the usage of cFDR beyond only GWAS to the accelerating field of functional genomics and can be applied iteratively to incorporate additional layers of data. We show through detailed simulations that Flexible cFDR increases sensitivity whilst controlling the FDR, and performs as well or better than the existing cFDR framework in the subset of use-cases supported by the latter.

We demonstrate the utility of our method by leveraging a variety of functional genomic data with GWAS *p*-values for asthma ^42^ to prioritise new genetic associations. We compare results to those from four existing methods which have previously been shown to outperform other approaches ^43,15^: a Bayesian approach, GenoWAP ^14^, grouping-based approaches, IHW ^6^ and FINDOR ^15^, and Boca and Leek’s FDR regression ^30^. We evaluate results according to validation status in the larger, independent, UK Biobank data set ^44^.

## Materials and methods

### Conditional false discovery rate

We begin by restating the definition and empirical estimator of the conditional Bayesian false discovery rate (cFDR). Consider *p*-values for *m* SNPs, denoted by *p*_1_*, …, p_m_*, corresponding to the null hypotheses of no association between the SNP and a principal trait (denoted by 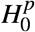). Let *p*_1_*, …, p_m_* be realisations from the random variable *P*. The Bayesian false discovery rate (FDR) is defined as the probability that the null hypothesis is true for a random SNP in a set of SNPs with *P* ≤ *p*:

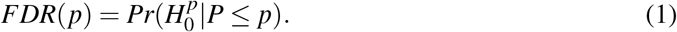

This Bayesian definition of a tail area FDR ^45^ is asymptotically equivalent ^46^ to the FDR introduced by Benjamini and Hochberg ^23^, which is the expected fraction of false discoveries amongst all discoveries.

Given additional *p*-values, *q*_1_*, …, q_m_*, for the same *m* SNPs for a “conditional trait”, the Bayesian FDR can be extended to the conditional Bayesian FDR (cFDR) by conditioning on both the principal and the conditional trait variables (in contrast to the standard FDR which conditions only on the principal trait variable). Assuming that *p_i_* and *q_i_* (for *i* = 1*, …, m*) are independent and identically distributed (iid) realisations of random variables *P, Q* satisfying:

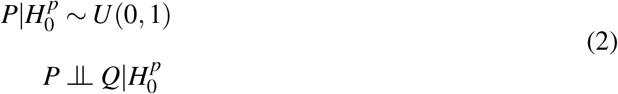

then the cFDR is defined as the probability 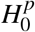 is true at a random SNP given that the observed *p*-values at that SNP are less than or equal to *p* in the principal trait and *q* in the conditional trait ^38^. Using Bayes theorem,

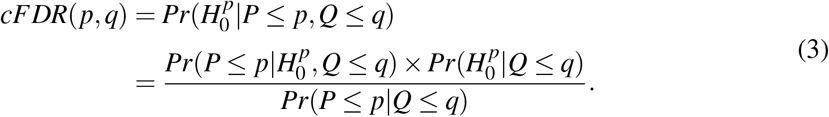

The cFDR framework implicitly assumes that there is a “positive stochastic monotonic relationship” between *p* and *q*, meaning that on average SNPs with smaller *p*-values in the conditional trait are enriched for smaller *p*-values in the principal trait. This assumption is naturally satisfied in the typical use-case of cFDR that leverages *p*-values for genetically related traits.

Using Bayes theorem and standard conditional probability rules, Eq (3) can be simplified to:

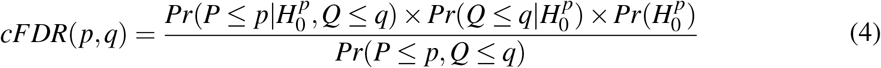

^47^.

It is conventional in the cFDR literature to conservatively approximate 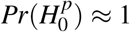, and this is reasonable in the GWAS setting where the proportion of true signals is expected to be very low (this may be debateable as sample sizes increase, but it is still appropriate in terms of being conservative). Given the assumptions in property (2), we can also approximate 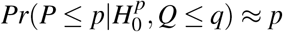, noting that this is an equality if *p* is correctly calibrated. The estimated cFDR is therefore:

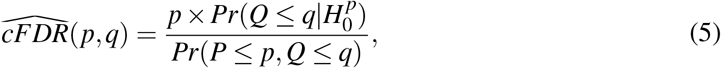

and existing methods use empirical cumulative distribution functions (CDFs) to estimate 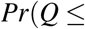 ^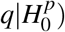^and *Pr*(*P* ≤ *p, Q* ≤ *q*) ^38,41^.

Having derived 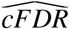 values for each *p*-value-covariate pair, a simple rejection rule would be to reject 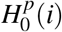 for any 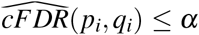, for 0 < *α* < 1. However, as discussed in Liley and Wallace ^41^, 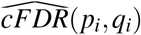 is not monotonically increasing with *p_i_* and we do not wish to reject the null for some (*p_i_, q_i_*) but not for some other pair (*p_j_, q _j_*) with *q_i_* = *q _j_* but *p _j_ < p_i_*.

Andreassen et al. ^38^ use the decision rule:

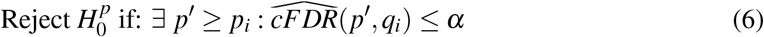

which closely follows the BH procedure ^23^. Yet unlike the BH procedure, this rejection rule does not control frequentist FDR at *α* ^47^. Liley and Wallace ^41^ described a method to control the frequentist FDR, but it is currently only suited to instances where the auxiliary data may be modelled using a mixture of centered normal distributions (for example by transforming auxiliary *p*-values to *Z* scores; 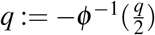.

### Flexible cFDR

The cFDR estimator in Eq (5) holds in the more general setting where *q*_1_*, …, q_m_* are real continuous values from some arbitrary distribution that is positively stochastically monotonic in *p*. However, the current methods to estimate the cFDR use empirical CDF estimates which can be inaccurate in regions of sparse data. These sparse data regions are likely to be found more often in unbounded auxiliary data from arbitrary distributions (for example near the extreme observations) than auxiliary data that are *p*-values from related traits, and thus bounded by [0, 1]. Moreover, the method used to control the frequentist FDR ^41^ requires normally distributed auxiliary data. We describe a new, more versatile cFDR framework for data pairs consisting of *p*-values for the principal trait (*p*) and continuous covariates from more general distributions (*q*). We call our method “Flexible cFDR”.

#### Flexible cFDR estimator

To estimate both 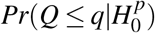 and *Pr*(*P* ≤ *p, Q* ≤ *q*) in Eq (5) we first fit a bivariate kernel density estimate (KDE) using a normal kernel. To do this, we begin by transforming the *p*-values for the principal trait (derived from a two-tailed test, as is typical in GWAS) to absolute *Z*-scores (*Z_p_*; since the sign of the associated *Z*-scores are essential arbitrary as they depend on which allele is designated “effect’). To avoid boundary effects which would otherwise bias estimates near 0, we mirror the absolute *Z*-scores onto the negative real line together with their associated *Q* values but only estimate the KDE on the non-negative part of the dat. We consequently model the PDF corresponding to *Z_p_, Q* in the usual way as

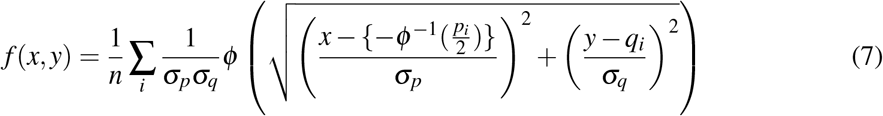

where *ϕ* is the standard normal density and the values *σ_p_* and *σ_q_* are the bandwidths determined using a well-supported rule-of-thumb ^48^, which assumes independent samples. Consequently, we fit the KDE to a subset of independent SNPs in the data set (independent SNP sets can be readily found using a variety of software packages including LDAK ^49^ and PLINK ^50^). We sum over *P* and *Q* to estimate *Pr*(*P* ≤ *p, Q* ≤ *q*).

Hard thresholding is used to approximate the distribution of 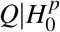 by *Q*|*P* > 1*/*2 in the earlier cFDR methods ^38,41^. Instead, in Flexible cFDR we empirically evaluate the influence of specific *p*-value quantities on the null hypothesis by utilising the local FDR framework, which estimates 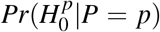 ^45^. We approximate 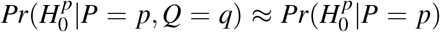 assuming that the majority of information about 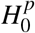is contained in *P* so that

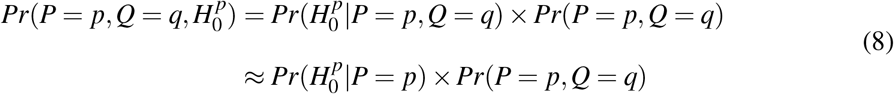

where *Pr*(*P* = *p, Q* = *q*) is estimated from our bivariate KDE and 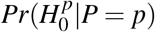 is estimated using the local FDR. Similarly to our method used to fit the KDE, to estimate the local FDR we mirror the absolute *Z*-scores onto the negative real line and extract the local FDR values for the non-negative part of the data, utilising the `locfdr R` package to do this.

From this, 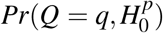 is estimated by integrating over *P* and 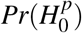 is then estimated by integrating over *Q* to obtain

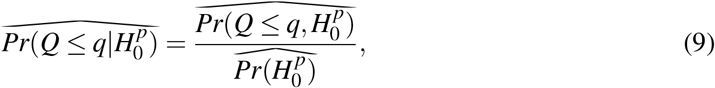

where we use ^ to denote that these are estimates under the assumption 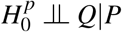.

Our final cFDR estimator is therefore:

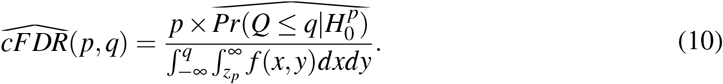

where *z_p_* is the *Z*-score associated with *p*.

As in the conventional cFDR approach, our estimator implicitly assumes a positive stochastic monotonic relationship between *p* and *q*. However, this is not guaranteed for the more general covariates that can now be leveraged with Flexible cFDR. If instead this relationship is negative (such that low *p*-values are enriched for high values of *q*), then the sign of the auxiliary data values can simply be reversed and the method can proceed as usual.

#### Mapping p-value-covariate pairs to v-values

We describe a similar approach to that in Liley and Wallace ^41^ but remove the restrictive parametric assumptions placed on the auxiliary data.

Following Liley and Wallace ^41^, we define “L-regions” as the set of points with 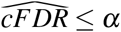 and the “L-curve” as the rightmost border of the L-region. Specifically, we calculate 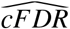 values for *p, q* pairs defined using a two-dimensional grid of *p* and *q* values. For each observed *p_i_, q_i_* pair we find the L-curve, which corresponds to the contour of estimated 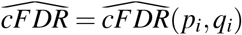. We then define the L-region from this L-curve.

We derive *v*-values, which are essentially the probability of a newly-sampled realisation (*p, q*) of *P, Q* falling in the L-region under 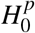. These are readily calculable by integrating the PDF of 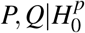, denoted by *f*_0_(*p, q*), over the L-region:

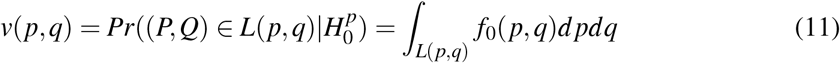

^41^. In the original method, *f*_0_(*p, q*) is estimated using a mixture-Gaussian distribution, but to support auxiliary data from arbitrary distributions (where the only distributional constraint is that it is positively stochastically monotonic in *p*) we utilise the assumptions in Eq (2) to write 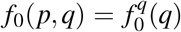 (since the PDF of p conditional on 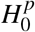 is the standard uniform density). We estimate 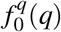 as an intermediate step in the derivation of 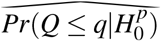.

The *v*-value can be interpreted as the probability that a randomly-chosen (*p, q*) pair has an equal or more extreme 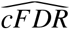 value than 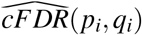 under 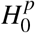 and is thus analogous to a *p*-value. We refer readers to Theorem 3.1 and its accompanying proof in Liley and Wallace (2021) ^41^ which shows that the *v*-values are uniformly distributed under the null hypothesis for *X* = (*p_i_, q_i_*) ∈ [0, 1]^2^, and this naturally holds for Flexible cFDR where *X* = (*p_i_, q_i_*) ∈ [0, 1] × [*q_low_, q_high_*] (where *q_low_* and *q_high_* are the lower and upper limits of the KDE support respectively).

Deriving *v*-values, which are analogous to *p*-values, means that the output from Flexible cFDR can be used directly in any conventional error rate controlling procedure, such as the BH method ^23^. The derivation of *v*-values also allow for iterative usage, whereby the *v*-values from the previous iteration are used as the “principal trait” *p*-values in the current iteration ^41^, thus allowing users to incorporate additional layers of auxiliary data into the analysis at each iteration, akin to leveraging multi-dimensional covariates.

#### Adapting to sparse data regions

To ensure that the integral of the KDE approximated in our method equals 1, we define the limits of its support to be 10% wider than the range of the data. This however introduces a sparsity problem whereby the data required to fit the KDE in or near these regions is very sparse. Adaptive KDE methods that find larger value bandwidths for these sparser regions are computationally impractical for large GWAS data sets. Instead, we opt to use left-censoring whereby all *q < q_low_* are set equal to *q_low_* and the value for *q_low_* is found by considering the number of data points required in a grid space to reliably estimate the density (S1 Fig). Note that since our method utilises cumulative densities, the sparsity of data for extremely large *q* is not an issue.

Occasionally, in regions where (*p, q*) are jointly sparse, the *v*-value can appear extreme compared to the *p*-value. To avoid artifactually inflating evidence for association, we fit a spline to *log*10(*v/p*) against *q* and calculate the distance between each data point and the fitted spline, mapping the small number of outlying points back to the spline and recalculating the corresponding *v*-value as required (S2 Fig).

#### Flexible cFDR software

We have created an R package, `fcfdr`, that implements the Flexible cFDR method. The software web-page (https://annahutch.github.io/fcfdr/) contains fully reproducible vignettes which illustrate how the Flexible cFDR method can be used to generate *v*-values from GWAS *p*-values and covariate data, and how these can be used directly in any error rate controlling procedure (for example using the `p.adjust` function with `method=“BH”` for FDR-adjusted *p*-values). We also include vignettes describing the types of auxiliary data that can be leveraged with Flexible cFDR, and how the LDAK method ^49^ can be used to generate an independent subset of SNPs for input into the software.

### Simulations

We used simulations to assess the performance of Flexible cFDR when iteratively leveraging various types of auxiliary data. We validated Flexible cFDR against the existing framework, which we call “empirical cFDR” ^41^, in two cases where *q* ∈ [0, 1] (as required by empirical cFDR). We then evaluated the performance of Flexible cFDR in three novel use-cases where the auxiliary data is no longer restricted to [0, 1]. We also analysed the simulation data using Boca and Leek’s FDR regression (referred to as BL) ^30^, which was the only other method that allowed for multiple covariates of this nature.

#### Simulating GWAS results (*p*)

We first simulated GWAS *p*-values for the arbitrary “principal trait” to be used as *p* in our simulations. We collected haplotype data from the UK10K project (REL-2012-06-02) ^51^ for 80,356 SNPs with minor allele frequency (MAF) ≥ 0.05 residing on chromosome 22. We split the haplotype data into 24 LD blocks representing approximately independent genomic regions defined by the LD detect method ^52^. Within each LD block, we sampled 2, 3 or 4 causal variants with log odds ratio (OR) effect sizes simulated from the standard Gaussian prior used for case-control genetic fine-mapping studies, *N*(0, 0.2^2^) ^53^ (the mean number of simulated causal variants in each simulation was 54). We then used the `simGWAS R` package ^54^ to simulate *Z*-scores from a GWAS in each region for 20,000 cases and 20,000 controls. We collated the *Z*-scores from each region and converted these to *p*-values representing the evidence of association between the SNPs and the arbitrary principal trait.

To generate an independent subset of SNPs required to fit the KDE, we converted the haplotype data to genotype data and used the `write.plink` function ^50^ to generate the files required for the LDAK software ^49^. We generated LDAK weights for each of the 80,356 SNPs and used the subset of 14,234 SNPs with non-zero LDAK weights as the independent subset of SNPs (an LDAK weight of zero means that its signal is (almost) perfectly captured by neighbouring SNPs) ^55^. Over the restricted interval of MAF values considered (MAF ≥ 0.05), we found that the MAF distributions of the whole SNP set and the independent subset were largely comparable, so we did not here perform the MAF matching procedure discussed below in our analysis of asthma data.

#### Simulating auxiliary data (q)

We consider five use-cases of cFDR (simulations A-E), defined by (i) the distribution of the auxiliary data *q* (ii) the relationship between *p* and *q* and (iii) the relationship between different *q* in each iteration (5 realisations of *q* were sampled in each simulation representing multi-dimensional covariates so that cFDR could be applied iteratively) (Table 1). We denote the value of *q* at SNP *i* in realisation *k* as 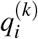.

**Table 1:** Summary of simulation analysis.

| Simulation | Distribution of $q$ | Relationship between $p_i$ and $q_i^{(k)}$ | Average pairwise correlation<sup>a</sup> between $q^{(k)}$ and $q^{(l)}$ |
| --- | --- | --- | --- |
| A | $p$-values (all null) | Independent | 0 |
| B | $p$-values (related trait) | Shared causal variants | 0.08 |
| C | Functional | Independent | 0 |
| D | Functional | Dependent | 0.08 |
| E | Functional | Dependent | 0.19 |

<sup>a</sup> Pairwise correlation values are the Pearson correlation coefficients.

In simulation A, we sampled *q_i_* ∼ *Uni f* (0, 1) to represent iterating over null *p*-values (S3A Fig). In simulation B, we investigated the standard use-case of cFDR by iterating over *p*-values from “related traits” (S3B Fig). To do this, we used the `simGWAS` R package ^54^ to simulate *p*-values, specifying the shared causal variants such that each pair of vectors *p, q*^(*k*)^ were *guaranteed* to share causal variants in exactly 4 of the 24 LD blocks, whilst each pair of vectors *q*^(*k*)^*, q*^(*j*)^ were *expected* to share causal variants in 4 of the 24 LD blocks.

In simulations C-E, we simulated auxiliary data that was no longer restricted to [0, 1]. In simulation C, we sampled *q_i_* from a bimodal mixture normal distribution that was independent of *p_i_*: *q_i_* ∼ 0.5 × *N*(−2, 0.5^2^) + 0.5 × *N*(3, 2^2^) (S3C Fig). In simulations D and E we simulated continuous auxiliary data that was dependent on *p_i_* by first defining “functional SNPs” as causal variants plus any SNPs within 10,000 bp, and “non-functional SNPs” as the remainder. In simulation D, we then sampled *q_i_* from different mixture normal distributions for functional and non-functional SNPs:

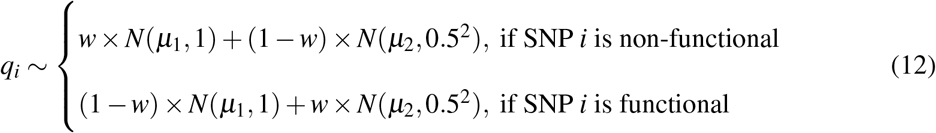

where *μ*_1_ ∈ {2.5, 3, 4}, *μ*_2_ ∈ {−1.5, −2, −3}, *w* ∈ {0.6, 0.7, 0.8, 0.9, 0.95} vary across iterations.

Since we anticipate our method being used to leverage functional genomic data iteratively, we also evaluated the impact of repeatedly iterating over auxiliary data that captures the same functional mark. To do this, in simulation E we iterated over realisations of *q* that are sampled from the same distribution,

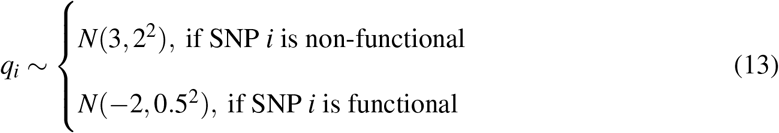

#### Running empirical cFDR, Flexible cFDR and BL

Following the vignette for the empirical cFDR software (https://github.com/jamesliley/cfdr/blob/master/vignettes/cfdr_vignette.Rmd), we first used the vl function to generate L-curves. As recommended in the documentation and to ensure that the rejection rules were not being applied to the same data from which they were determined, we used the leave-one-out-procedure whereby L-curves were fit separately for data points in each LD block using data points from the other LD blocks. To ensure that the cFDR curves were strictly decreasing (preventing a complication whereby all *v*-values corresponding to the smallest *p*-values were given the same value), we reduced the value of the `gx` parameter to the minimum *p*-value in the LD block. We then estimated the distribution of *P, Q*|*H*_0_ using the `fit.2g` function and integrated its density over the computed L-regions using the `il` function, specifying a mixture Gaussian distribution for the *Z*-scores.

Flexible cFDR was implemented using the `flexible_cfdr` function in the `fcfdr R` package with default parameter values. Both cFDR methods were applied iteratively 5 times in each simulation to represent leveraging multi-dimensional covariates.

For BL, we used the `lm_qvalue` function in the `swfdr` Bioconductor R package (version 1.16.0) to derive adjusted *p*-values. The covariate matrix that we used consisted of five columns for *q*1*, q*2*, q*3*, q*4*, q*5.

#### Evaluating sensitivity, specificity and FDR control

To quantify the results from our simulations, we used the BH procedure to derive FDR-adjusted *v*-values from empirical and Flexible cFDR (which we call “FDR values” for conciseness). For BL, we used the adjusted *p*-values as the quantity of interest. We then calculated proxies for the sensitivity (true positive rate) and the specificity (true negative rate) at an FDR threshold of *α* = 5*e* − 06, which roughly corresponds to the genome-wide significance *p*-value threshold of 5*e* − 08 (S4 Fig). We defined a subset of “truly associated SNPs” as any SNPs with *r*^2^ ≥ 0.8 with any of the causal variants. Similarly, we defined a subset of “truly not-associated SNPs” as any SNPs with *r*^2^ ≤ 0.01 with all of the causal variants. (Note that there are 3 non-overlapping sets of SNPs: “truly associated”, “truly not-associated” and neither of these). We calculated the sensitivity proxy as the proportion of truly associated SNPs that were called significant and the specificity proxy as the proportion of truly not-associated SNPs that were called not significant. To examine whether our results were robust to using different *r*^2^ values to define truly associated and truly not-associated SNPs, we also evaluated our sensitivity and specificity proxies using larger *r*^2^ values.

To assess whether the FDR was controlled within a manageable number of simulations, we raised *α* to 0.05 and calculated the proportion of SNPs called FDR significant which were truly not-associated (that is, *r*^2^ ≤ 0.01 with all of the causal variants).

### Application to asthma

We demonstrate the utility of our method by leveraging a variety of functional genomic data with GWAS *p*-values for asthma ^42^. Specifically, we describe two applications: (1) leveraging GenoCanyon scores with asthma GWAS *p*-values and (2) leveraging ChIP-seq data in relevant cell types with asthma GWAS *p*-values. We compare the performance of Flexible cFDR to that of four existing methods in the applications which support their usage.

#### Asthma GWAS data

Asthma GWAS summary statistics for 2,001,256 SNPs were downloaded from the NHGRI-EBI GWAS Catalog ^57^ for study accession GCST006862^42^ on 10/10/2019. We used the *p*-values generated from a meta-analysis of 56 GWASs for individuals of European ancestry under a random effects model, totalling 19,954 asthma cases and 107,715 controls. The genomic inflation factor for this study was *λ* = 1.055, implying minimal inflation of test statistics. The UCSC liftOver utility ^58^ was used to convert GRCh38/hg38 into GRCh37/hg19 coordinates, and those that could not be accurately converted were removed. All co-ordinates reported are for GRCh37/hg19. We call this GWAS data the “discovery GWAS data set”.

We analysed these data with methods that leverage auxiliary data as described below, and evaluated results using data from a larger asthma GWAS performed by the Neale Lab (self-reported asthma: 20002_1111) for 41,934 asthma cases and 319,207 controls from UK Biobank ^44^ (URL: https://www.dropbox.com/s/kp9bollwekaco0s/20002_1111.gwas.imputed_v3.both_sexes.tsv.bgz?dl=0 downloaded on 10/05/2020). Specifically, if a SNP was claimed to be significant in the discovery GWAS data set or after applying cFDR/ related methods, and it was also significant in the Neale Lab UK Biobank validation data, then we say that it is validated. We used the BH procedure to derive FDR-adjusted *p*-values (which we call “FDR values” for conciseness) and defined significant SNPs as those with *FDR* ≤ 0.000148249, which corresponds to the genome-wide significance *p*-value threshold of *p* ≤ 5*e* − 08 (0.000148249 is the maximum FDR value amongst SNPs with raw *p*-values ≤ 5*e* − 08 in the discovery GWAS data set). We restricted analysis to the 1,968,651 SNPs that were present in both the discovery and the validation GWAS data sets.

We down-sampled the independent subset of SNPs to match the MAF distribution in this subset to that in the whole set of SNPs. This accounts for the confounding of LDAK weights and GWAS *p*-values by MAF: less common SNPs (*MAF* < 0.05) are over-represented among the independent subset and have, on average, larger *p*-values. Matching in this way prevents a bias of the KDE fit towards the behaviour of rarer SNPs. We used MAFs estimated from the CEU sub population of the 1000 Genomes resource ^59^, and for the 639 SNPs with missing MAF values we used values randomly sampled from the empirical MAF distribution derived from the other SNPs. This reduced the independent subset of SNPs for fitting the KDE from 509,716 to 247,879 SNPs.

To identify independent hits we used the LD clumping algorithm in PLINK 1.9^50^, using a 5 Mb window and an *r*^2^ threshold of 0.01^15^. We used haplotype data from the 503 individuals of European ancestry from 1000 Genomes project Phase III ^59^ as a reference panel to calculate LD between SNPs. The SNP with the smallest *p*-value in the discovery GWAS data set in each LD clump was called the “index SNP”.

#### Application 1: Leveraging GenoCanyon scores

Tools have now been developed that integrate various genomic and epigenomic annotation data to quantify the pathogenicity, functionality and/or deleteriousness of both coding and non-coding GWAS variants ^60,61,62,63,64^. For example, GenoCanyon scores aim to infer the functional potential of each position in the human genome ^64^. They are derived from the union of 22 computational and experimental annotations (broadly falling into conservation measure, open chromatin, histone modification and TFBS categories) in different cell types. We downloaded GenoCanyon scores for each of the 1,968,651 SNPs that GWAS *p*-values were available for, noting a bimodal distribution for the scores (S5A Fig).

#### Application 2: Leveraging ChIP-seq data

The histone modification H3K27ac is associated with active enhancers ^65^ and so SNPs residing in genomic regions with high H3K27ac counts in trait-relevant cell types may be more likely to be associated with the trait of interest ^66^.

We downloaded consolidated fold-enrichment ratios of H3K27ac ChIP-seq counts relative to expected background counts from NIH Roadmap ^67^ in tissues relevant for asthma: immune cells and lung tissue. We mapped each SNP in our GWAS data set to its corresponding genomic region and recorded the H3K27ac fold change values for each SNP in each cell type using bedtools intersect ^68^. For SNPs on the boundary of a genomic region (and therefore mapping to two regions) we randomly selected one of the regions.

The raw H3K27ac fold change data had very long tails and so we transformed the values: *q* := *log*(*q* + 1). We observed that the data for the different cell types roughly fell into two clusters (S6 Fig): lymphoid cells (consisting of CD19^+^, CD8^+^ memory, CD4^+^ CD25^−^ CD45RA^+^ naive, CD4^+^ CD25^−^ CD45RO^+^ memory, CD4^+^ CD25^+^ CD127^−^ Treg, CD4^+^ CD25int CD127^+^ Tmem, CD8 naive, CD4 memory, CD4 naive and CD4^+^ CD25^−^ Th cells) clustered with CD56 cells whilst lung tissue clustered with monocytes (CD14^+^ cells). We therefore averaged the transformed H3K27ac fold change values in lymphoid and CD56 cell types to derive *q*1, and the transformed H3K27ac fold change values in lung tissue and monocytes to derive *q*2, adding a small amount of noise [*N*(0, 0.1^2^)] to the latter to smooth out the discrete valued counts (S7 Fig).

### Analysis methods

#### Flexible cFDR

We used the `flexible_cfdr` function in the `fcfdr` R package to generate *v*-values derived by leveraging the auxiliary data described above with asthma GWAS *p*-values. We defined our independent SNP set as the set of 509,716 SNPs given a non-zero LDAK weight ^55^ for the `indep_index` parameter and we used the optional `maf` parameter to supply the MAF values required for our MAF matching procedure. We used the BH procedure to derive FDR values and used these as the output of interest.

#### GenoWAP

GenoWAP is a Bayesian method that leverages GenoCanyon scores with GWAS *p*-values to output posterior scores of disease-specific functionality for each SNP ^14^. The GenoWAP software requires a `threshold` parameter defining functional SNPs according to their GenoCanyon score. For this, we used the default recommended value of 0.1, which corresponded to 40% of the SNPs in our data set being “functional”. We used the `GenoWAP.py` python script to obtain posterior scores for each SNP, and used these as the output of interest. GenoWAP could only be used for application 1 because it only supports auxiliary data that is GenoCanyon scores.

#### IHW

Independent hypothesis weighting (IHW) ^6^ is a statistical method for covariate-informed multiple testing whereby variables are divided into groups and optimal group specific weights are derived (which maximise the number of discoveries whilst controlling the FDR) for use in a weighted BH procedure. We used the `IHW` Bioconductor R package (version 1.18) with default parameters, specifying the level of FDR control `alpha=0.000148249`, and used the adjusted *p*-values as the output of interest. IHW could only be used for application 1 because it is not clear how to use this method to leverage multi-dimensional covariate vectors.

#### Boca and Leek’s FDR regression

BL estimates the proportion of null hypotheses conditional on observed covariates and uses these as plug-in estimates for the FDR ^30^. We used the `swfdr` Bioconductor R package (version 1.16.0) to derive adjusted *p*-values ^56^ and used these as the output of interest. For application 1, the `lm_qvalue` function was used with a covariate matrix consisting of a single column of GenoCanyon scores for each SNP. For application 2, the `lm_qvalue` function was used with a covariate matrix consisting of two columns for *q*1 and *q*2.

#### FINDOR

FINDOR is a *p*-value re-weighting method which leverages a wider range of non-cell-type-specific functional annotations. FINDOR uses the baseline-LD model from Gazal et al. ^27^ for prediction, and so we were unable to directly compare the methods when leveraging the same GenoCanyon or ChIP-seq auxiliary data. Instead, as recommended we used FINDOR to leverage the 96 annotations from the latest version of the baseline-LD model (version 2.2) with asthma GWAS *p*-values. Briefly, this auxiliary data contains the 75 annotations from Gazal et al. ^27^ (including functional regions, histone marks, MAF bins and LD-related annotations) plus extra annotations including synonymous/ non-synonymous, conserved annotations, 2 flanking bivalent TSS/ enhancer annotations from NIH Roadmap ^67^, promoter/ enhancer annotations ^69^, promoter/ enhancer sequence age annotations ^70^ and 11 new annotations from Hujoel et al. ^71^ (5 new binary annotations and corresponding flanking annotations and 1 continuous count annotation). We matched SNPs to their annotations using rsID and GRCh37/hg19 coordinates.

To run FINDOR, stratified LD score regression (S-LDSC) must first be implemented to obtain anno-456 tation effect size estimates, 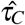. To run S-LDSC, we downloaded (i) partitioned LD baseline-LD model v2.2^27^, (ii) regression weight LD scores and (iii) allele frequencis for available variants in the 1000 Genomes Phase 3 data set. We then used the `munge_sumstats.py` python script in the `ldsc` package to convert the asthma GWAS summary statistics to the correct format for use in the `ldsc` software. We restrict analysis to HapMap3 SNPs using the `–merge-alleles` flag, as recommended in the LDSC and FINDOR documentation.

We ran S-LDSC with the `–print-coefficients` flag to generate the `.result` file containing the annotation effect size estimates required for FINDOR. Specifically, pre-computed regression weight LD scores were read in for 1,187,349 variants, for which 1,034,758 remained after merging with reference panel SNP LD scores, for which 1,032,395 SNPs remained after merging with regression SNP LD scores.

To run FINDOR, partitioned LD scores must also be supplied for the SNPs in the data set. To do this, we downloaded the 1000 Genomes EUR Phase 3 PLINK files and annotation data and followed the ‘LD Score Estimation Tutorial’ on the LDSC GitHub page. Partitioned LD scores could be generated for the 1,976,360 (out of 2,001,256) SNPs in the asthma data set that were also present in the 1000 Genomes Phase 3 data set.

We then generated a file for the asthma GWAS data ^42^, including columns for sample sizes, SNP IDs and *Z*-scores. We used this file, along with the computed partitioned LD scores and the `.result` file from S-LDSC to obtain re-weighted *p*-values for the 1,968,651 SNPs using FINDOR. We used the BH procedure to convert these to FDR values and used these as our output of interest.

## Results

### Simulations show Flexible cFDR controls FDR and increases sensitivity where appropriate

We expect that leveraging irrelevant data should not change our conclusions about a study. Simulations A and C showed that the sensitivity and specificity remained stable across iterations and that the FDR was controlled at a pre-defined level when leveraging independent auxiliary data with Flexible cFDR (Fig 1A, Fig 1C). In contrast, when leveraging relevant data, we hope that the sensitivity improves whilst the specificity remains high. This is what we observed for Flexible cFDR in simulations B and D (Fig 1B, Fig 1D). The increase in sensitivity related to how informative the auxiliary data was, whereby the sensitivity generally increased more in simulation B than simulation D, where the average Pearson correlation coefficient between *p* and *q*^(*k*)^ was *r* = 0.06 and *r* = 0.04 respectively. These findings were robust to the *r*^2^ threshold used to define our sensitivity (S8 Fig) and specificity (S9 Fig) proxies.

**Figure 1:**
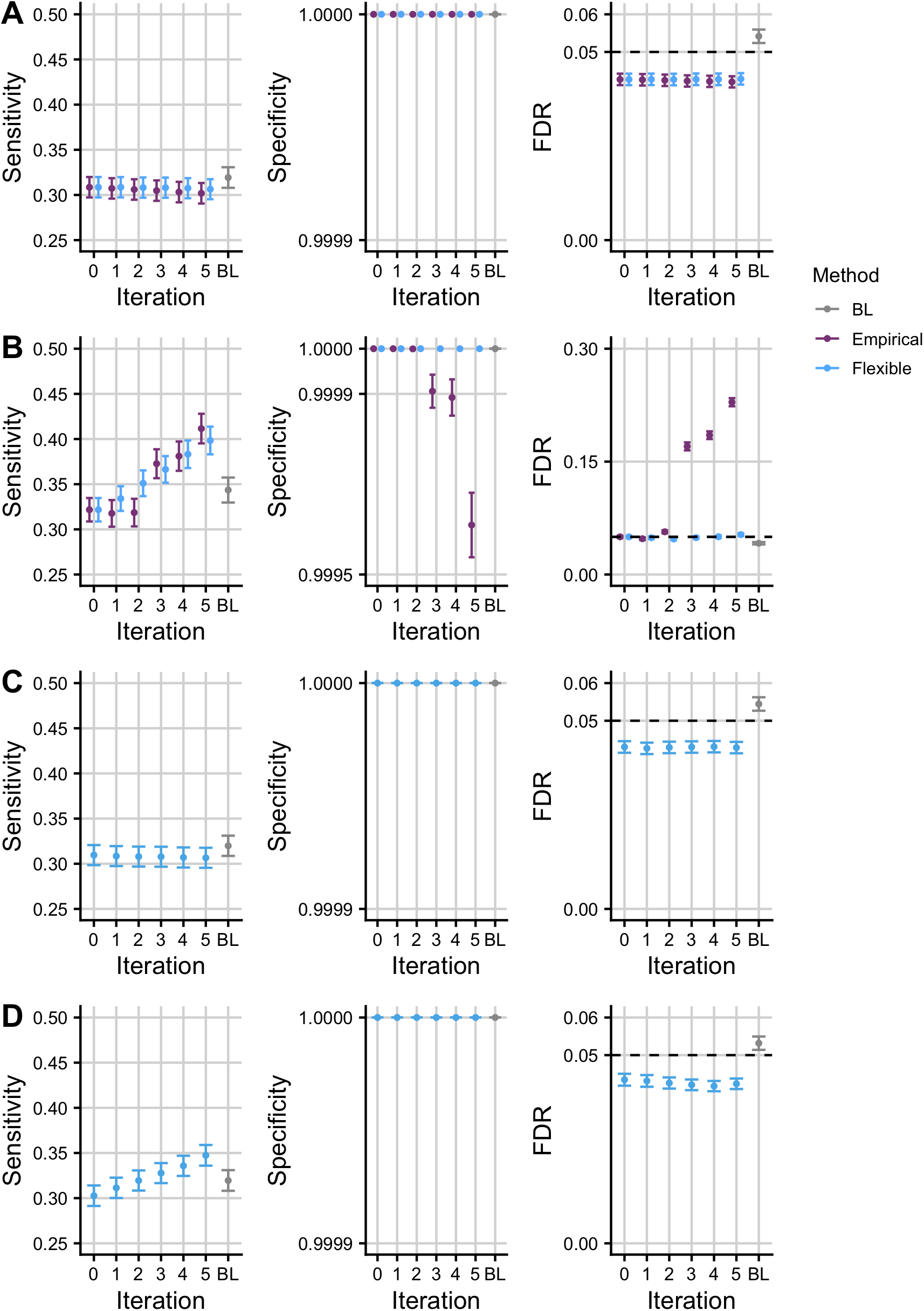
Simulation results. Mean +/− standard error for the sensitivity, specificity and FDR of FDR values from empirical and Flexible cFDR when iterating over independent (A; “simulation A”) and dependent (B; “simulation B”) auxiliary data that is bounded by [0, 1]. Panels C and D show the results from Flexible cFDR when iterating over independent (C; “simulation C”) and dependent (D; “simulation D”) auxiliary data simulated from bimodal mixture normal distributions. BL refers to results when using Boca and Leek’s FDR regression to leverage the 5-dimensional covariate data. Iteration 0 corresponds to the original FDR values. Our sensitivity proxy is calculated as the proportion of SNPs with *r*^2^ ≥ 0.8 with a causal variant (“truly associated”), that were detected with a FDR value less than 5*e* − 06. Our specificity proxy is calculated as the proportion of SNPs with *r*^2^ ≤ 0.01 with all the causal variants (“truly not-associated”), that were not detected with a FDR value less than 5*e* − 06. Our FDR proxy is calculated as the proportion of SNPs that were detected with a FDR value less than 0.05, that had *r*^2^ ≤ 0.01 with all the causal variants (“truly not-associated”) (we raised *α* to 0.05 in order to assess FDR control within a manageable number of simulations). Results were averaged across 100 simulations.

For simulations A and B, we could compare Flexible cFDR performance to that of the current method, empirical cFDR, since *q* ∈ [0, 1] ^41^. Performance was similar for simulation A, whilst for simulation B, the sensitivity of the two methods was comparable but empirical cFDR exhibited a greater decrease in the specificity and failed to control the FDR in later iterations (Fig 1B). This contrasts with earlier results for empirical cFDR, which showed good control of FDR ^41^, and reflects the structure of our simulations which assume dependence between different realisations of *q*.

A leave-one-out procedure is required for the empirical cFDR method, as it utilises empirical CDFs and including an observation when estimating its own L-curve causes the curve to deviate around the observed point ^41^. Flexible cFDR does not require a leave-one-out procedure as KDEs are used instead of empirical CDFs. Additionally, Flexible cFDR is quicker to run than empirical cFDR, taking approximately 3 minutes compared to empirical cFDR which takes approximately 6 minutes to complete a single iteration on 80,356 SNPs (using one core of an Intel Xeon E5-2670 processor running at 2.6GHz). Together, these findings indicate that Flexible cFDR performs no worse, and generally better, than empirical cFDR in use-cases where both methods are supported.

We also benchmarked the performance of Flexible cFDR against that of BL ^30^, which was the only other method that allows for multiple covariates of this nature. We found that BL failed to control the FDR when leveraging independent covariate data, which may be due to the correlations between SNPs (Fig 1A, Fig 1C). Indeed, Boca and Leek ^30^ found that control of the FDR by BL was worse with increasing correlation, but we note that correlation structure is ubiquitous in GWAS data. When leveraging dependent covariate data, BL was consistently less powerful than Flexible cFDR (Fig 1B, Fig 1D) and it failed to control the FDR in simulations leveraging dependent covariates from arbitrary distributions (Fig 1D), which represent the general use-case of the method ^30^.

We anticipate that Flexible cFDR will typically be used to leverage functional genomic data iteratively and it is helpful that specificity remains high and FDR is controlled in simulation D. It is obvious that repeated conditioning on the *same* data would produce erroneous results, with SNPs with a modest *p* but extreme *q* incorrectly attaining greater significance with each iteration. For strict validity, we require 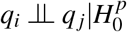 as the *v*-value from iteration *i* will contain some information about *q_i_*, and the cFDR assumes 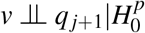 at the next iteration. However, even when 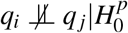, we expect the dependence between *v* and *q* to be quite weak, hence the acceptable FDR control in simulations B and D above.

Given the wealth of functional marks available for similar tissues and cell types (for example subsets of peripheral immune cells), we wanted to assess robustness of our procedure to more extreme dependence by repeatedly iterating over auxiliary data that is capturing the same functional mark. In simulation E, the sensitivity increased with each iteration at the expense of a drop in the specificity and loss of FDR control in later iterations (Fig). The drop in specificity and loss of FDR control is exacerbated when iterating over *exactly* the same auxiliary data in each iteration (S10 Fig), as expected. We therefore recommend that care should be taken not to repeatedly iterate over functional data that is capturing the same genomic feature, and in a real data example that follows, we average over cell types which show correlated values for functional data.

### Analysis of asthma GWAS leveraging GenoCanyon scores

Overall, 655 SNPs were FDR significant (*FDR* ≤ 0.000148249) in the original asthma GWAS ^42^. We used Flexible cFDR to leverage GenoCanyon scores measuring SNP functionality with asthma GWAS *p*-values (S1 Table). SNPs with high GenoCanyon scores were enriched for smaller asthma *p*-values (S11 Fig) and accordingly FDR values from Flexible cFDR for SNPs with high GenoCanyon scores (and therefore more likely to be functional) were lower than their corresponding original FDR values, whilst those for SNPs with low GenoCanyon scores (and therefore less likely to be functional) were higher than their corresponding original FDR values (S12 Fig). (Due to the positive stochastic monotonicity requirement for cFDR, the Flexible cFDR software reversed the sign of the GenoCanyon scores for its internal calculations). Specifically, Flexible cFDR identified 12 newly FDR significant SNPs (rs4705950, rs6903823, rs9262141, rs1264349, rs2106074, rs3130932, rs9268831, rs3129719, rs1871665, rs16924428, rs1663687 and rs12900122) which had high GenoCanyon scores (mean GenoCanyon score = 0.77) and 3 SNPs were no longer FDR significant which had low GenoCanyon scores (mean GenoCanyon score = 0.01). At the locus level no newly significant, or newly not-significant, loci were identified.

We compared the results from Flexible cFDR to those from IHW ^6^, BL ^30^ and GenoWAP ^14^ when leveraging the exact same auxiliary data (S1 Table). IHW groups SNPs based on their covariate values and derives optimal group-specific weights for use in a weighted BH procedure. Interestingly, all SNPs were allocated a weight of 1 in this instance, meaning that IHW reduced to the conventional BH procedure (and so the adjusted *p*-values from IHW were identical to the original FDR values from Demenais et al. ^42^).

In BL, logistic regression is used to estimate how the distribution of input *p*-values depends on the GenoCanyon scores to estimate the probability that the null hypothesis of no association is true for each SNP. These probabilities ranged from 0.957 for the SNP with the largest GenoCanyon score to 0.993 for the SNP with the smallest GenoCanyon score (S5B Fig). The consequence of the narrow range of these values is that the adjusted values from BL were very similar to the original FDR values (S13 Fig). Specifically, BL only identified 3 newly FDR significant SNPs, and these were all also identified by Flexible cFDR. One of these had a very high GenoCanyon score (rs1871665 with score = 0.999) whilst the other two had medium (rs16924428 with score = 0.532) or low (rs9268831 with score = 0.224) scores. No SNPs were identified as no longer FDR significant after applying BL and at the locus level, no newly significant, or newly not-significant, loci were identified.

Since GenoWAP outputs posterior probabilities rather than *p*-values, we compared the performance of the methods with GenoWAP based on the rankings of SNPs using the UK Biobank data resource. Firstly, at the SNP-level, for each of the 5152 SNPs that passed FDR significance in the UK Biobank data, we compared the rank of the FDR value in the discovery data set with (1) the rank of the FDR value after applying Flexible cFDR, (2) the rank of the FDR value from BL and (3) the rank of the (negative) posterior score from GenoWAP. (IHW was not included in this comparison because the output from IHW was just the original FDR values). We found that the percentage of FDR significant SNPs in the UK Biobank data which had an improved rank after applying each of the methods was similar (Table 2) and that 68.5% of the SNPs that improved ranks in at least one of the methods improved rank in all of the methods. Similarly, the percentage of the 1,963,499 SNPs that were not FDR significant in UK Biobank which had a decreased rank after applying each of the methods was similar (Table 2) and 49.6% of the SNPs that decreased rank in at least one of the methods decreased rank in all of the methods.

**Table 2:** Summary of SNP-level results when leveraging GenoCanyon scores with asthma GWAS *p*-values. Table lists the percentage of the 5152 FDR significant SNPs in UK Biobank which improved rank (“UK Biobank significant which improved rank”) and the percentage of the 1,963,499 SNPs that were not FDR significant in UK Biobank which decreased rank (“UK Biobank not-significant which decreased rank”) after applying Flexible cFDR, BL or GenoWAP.

|  | Flexible cFDR | BL | GenoWAP |
| --- | --- | --- | --- |
| UK Biobank significant which improved rank | 60.3% | 52.4% | 61.5% |
| UK Biobank not-significant which decreased rank | 46.8% | 58.0% | 44.8% |

Secondly, we focused on the 114 loci that passed FDR significance in the UK Biobank data set. Within each of the 114 loci, we identified the SNP with the lowest *p*-value and called this the “index SNP”. For each index SNP, we compared the rank of the FDR values in the discovery GWAS data set with (1) the rank of the FDR value after applying Flexible cFDR, (2) the rank of the FDR value from BL and (3) the rank of the (negative) posterior score from GenoWAP. In total, 42.1% had an improved rank after applying Flexible cFDR compared to 28.9% after BL and 55.3% after GenoWAP (and 70.1% of these were shared by all three methods) (Table 3). Similarly, of the 301 loci that were not FDR significant in UK Biobank, 40.5% had a decreased rank after applying Flexible cFDR compared with 40.9% after BL and 34.6% after GenoWAP (and 51.4% of these were shared by all three methods) (Table 3).

**Table 3:** Summary of locus-level results when leveraging GenoCanyon scores with asthma GWAS *p*-values. Table lists the percentage of the 114 FDR significant index SNPs in UK Biobank which improved rank (“UK Biobank significant which improved rank”) and the percentage of the 301 index SNPs that were not FDR significant in UK Biobank which decreased rank (“UK Biobank not-significant which decreased rank”) after applying Flexible cFDR, BL or GenoWAP.

|  | Flexible cFDR | BL | GenoWAP |
| --- | --- | --- | --- |
| UK Biobank significant which improved rank | 42.1% | 28.9% | 55.3% |
| UK Biobank not-significant which decreased rank | 40.5% | 40.9% | 34.6% |

In all, the results were similar for Flexible cFDR, BL and GenoWAP when leveraging GenoCanyon scores of SNP functionality with asthma GWAS *p*-values, but rather unexciting as no newly significant loci were identified. We suggest that this is due to the one-dimensional non-trait-specific auxiliary data that is being leveraged, which is unlikely to capture enough disease relevant information to substantially alter conclusions from a study. This is supported by our intermediary results, where the optimal weights derived in IHW were all equal to 1 and the estimated proportions of true null hypotheses conditional on the GenoCanyon scores in BL were almost negligible.

### Analysis of asthma GWAS leveraging ChIP-seq data uncovers new genetic associations

In agreement with reports that GWAS SNPs are enriched in active chromatin ^72^, we observed that H3K27ac fold change values in asthma relevant cell types were negatively correlated with asthma GWAS *p*-values (S14 Fig) such that SNPs with high fold change values were enriched for smaller *p*-values (S15 Fig). (Due to the positive stochastic monotonicity requirement for cFDR, the Flexible cFDR software reversed the sign of the fold change values for its internal calculations). Accordingly, FDR values from Flexible cFDR for SNPs with high H3K27ac fold-change counts in asthma relevant cell types were lower than their corresponding original FDR values, whilst those for SNPs with low H3K27ac fold-change counts in asthma relevant cell types were higher than their corresponding original FDR values (Fig 3A; Fig 3B; Fig 3C; S2 Table).

**Figure 2:**
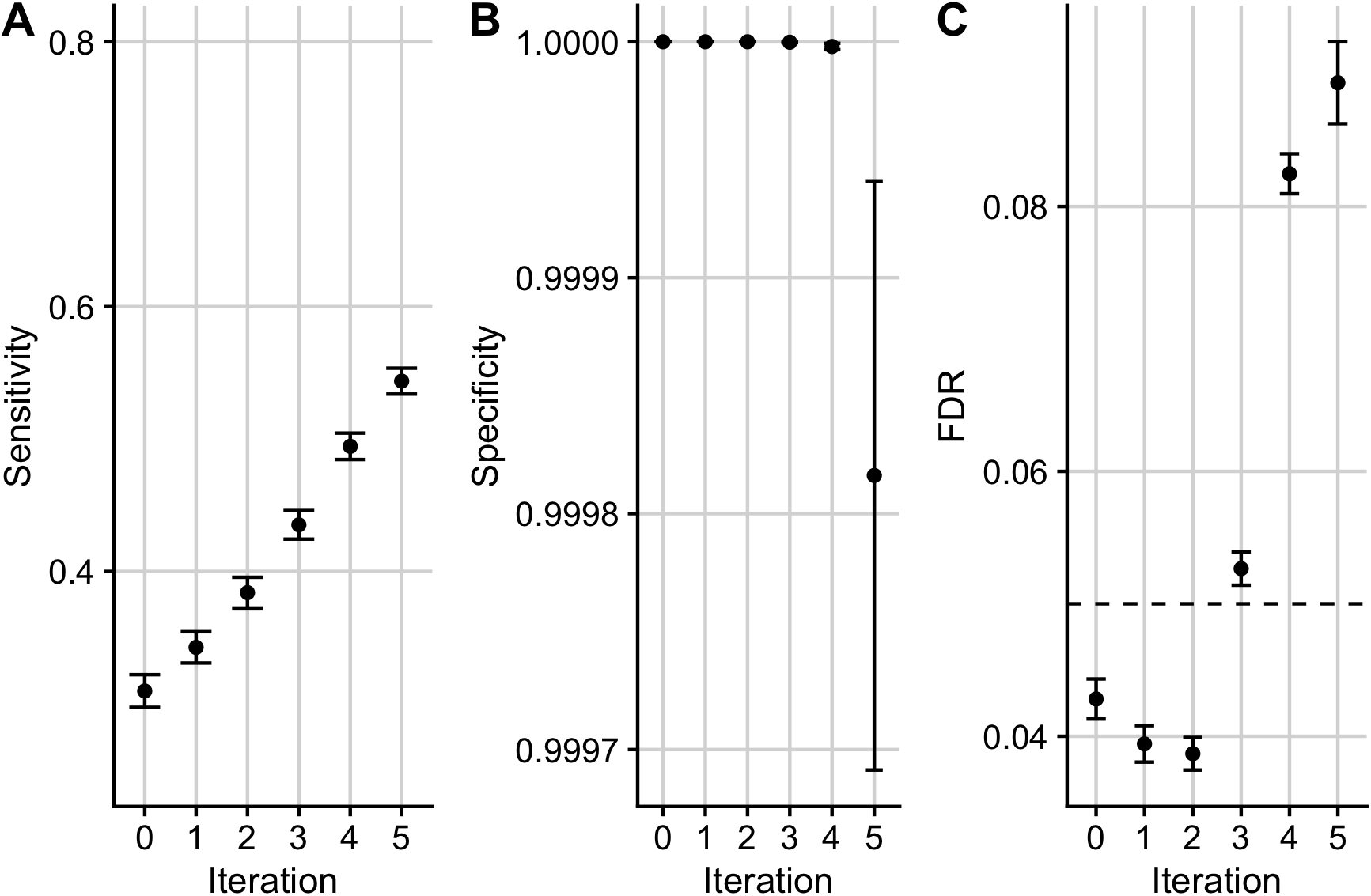
Simulation results for scenario E. Mean +/− standard error for the sensitivity (A), specificity (B) and FDR (C) of FDR values from Flexible cFDR when iterating over auxiliary data sampled from the same distribution (“simulation E”). Iteration 0 corresponds to the original FDR values. Our sensitivity proxy is calculated as the proportion of SNPs with *r*^2^ ≥ 0.8 with a causal variant (“truly associated”), that were detected with a FDR value less than 5*e* − 06. Our specificity proxy is calculated as the proportion of SNPs with *r*^2^ ≤ 0.01 with all the causal variants (“truly not-associated”), that were not detected with a FDR value less than 5*e* − 06. Our FDR proxy is calculated as the proportion of SNPs that were detected with a FDR value less than 0.05, that had *r*^2^ ≤ 0.01 with all the causal variants (“truly not-associated”) (we raised *α* to 0.05 in order to assess FDR control within a manageable number of simulations). Results were averaged across 100 simulations.

**Figure 3:**
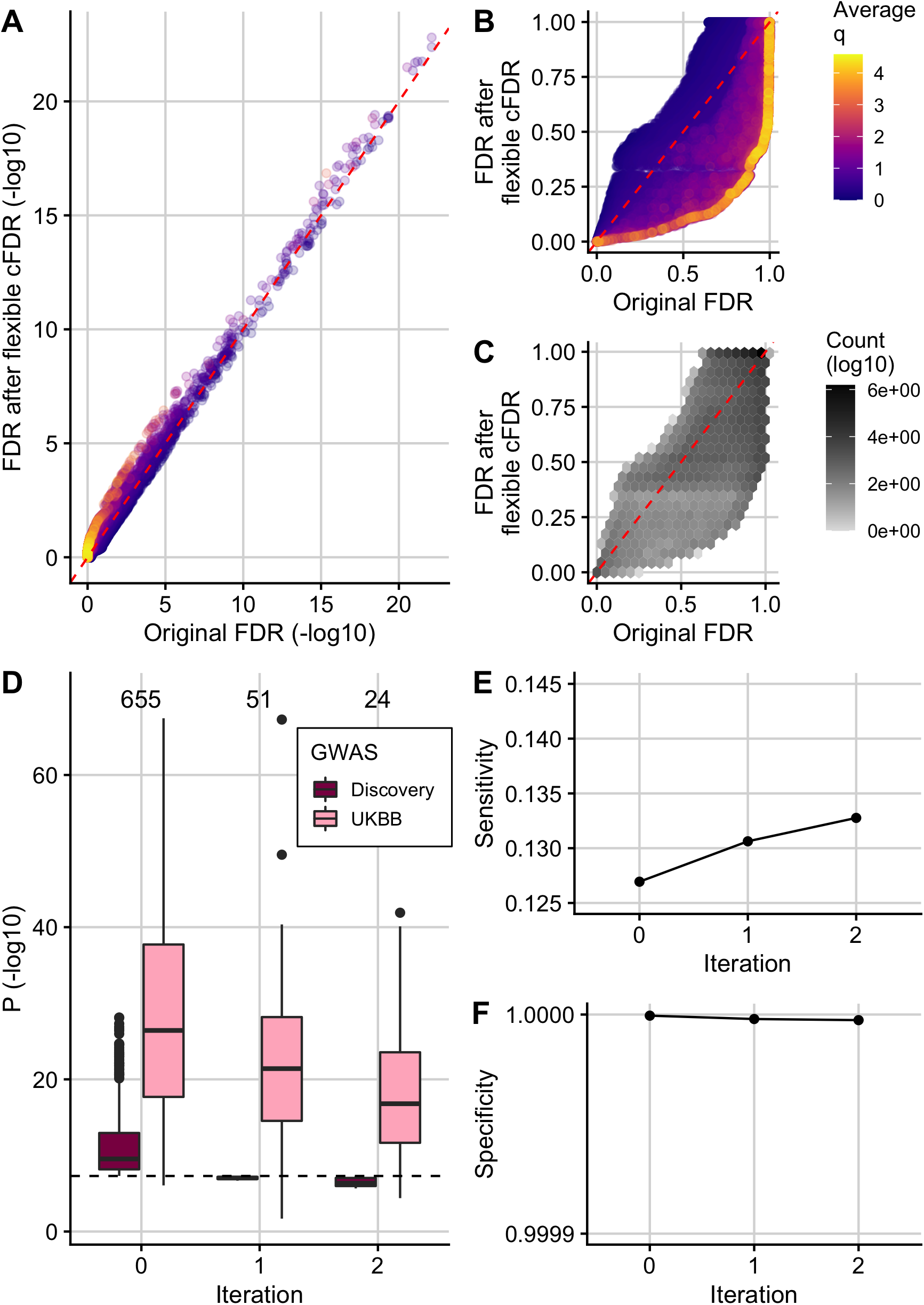
Using Flexible cFDR to leverage H3K27ac data with asthma GWAS *p*-values. (A) (-log10) FDR values after 2 iterations of Flexible cFDR leveraging H3K27ac counts in relevant cell types against raw (-log10) FDR values coloured by the average value of the auxiliary data across iterations. (B) As in A but non-log-transformed FDR values. (C) As in B but coloured by (log10) counts of data points in each hexbin. (D) Box plots of (-log10) *p*-values in the discovery GWAS and the UK Biobank data set for the 655 SNPs that were FDR significant in the original GWAS (Iteration 0), 51 SNPs that were newly FDR significant after iteration 1 of Flexible cFDR (leveraging average H3K27ac fold change values in lymphoid and CD56 cell types) and 24 SNPs that were newly FDR significant after iteration 2 of Flexible cFDR (subsequently leveraging average H3K27ac fold change values in lung tissue and CD14^+^ cells). Black dashed line at genome-wide significance (*p* = 5*e* − 08). (E) Sensitivity proxy and (F) specificity proxy for the H3K27ac application results. Sensitivity proxy is calculated as the proportion of SNPs that are FDR significant in the UK Biobank data set that are also FDR significant in the original GWAS (iteration 0), after iteration 1 of Flexible cFDR or after iteration 2 of Flexible cFDR. Specificity is calculated as the proportion of SNPs that are not FDR significant in the UK Biobank data set that are also not FDR significant in the original GWAS (iteration 0), after iteration 1 of Flexible cFDR or after iteration 2 of Flexible cFDR.

The 655 SNPs that were FDR significant (*FDR* ≤ 0.000148249) in the original asthma GWAS ^42^ have strong replication *p*-values in the UK Biobank data set used for validation (Fig 3D; Iteration 0). By leveraging H3K27ac data, Flexible cFDR identified weaker signals that were not significant in the original data but have reassuringly small *p*-values in the UK Biobank data (median *p*-value in UK Biobank data for these SNPs is 4.65*e* − 21; Fig 3D). Specifically, Flexible cFDR identified 51 newly significant SNPs when leveraging average H3K27ac fold change values in lymphoid and CD56 cell types (Fig 3D; Iteration 1), and 24 newly significant SNPs when subsequently leveraging average H3K27ac fold change values in lung tissue and monocytes (Fig 3D; Iteration 2). The maximum *p*-value for the 69 newly significant SNPs (6 SNPs newly significant after iteration 1 were no longer significant after iteration 2) in the discovery GWAS data set was 2.15*e* − 06 and the maximum UK Biobank *p*-value for these SNPs was 0.02. The newly significant SNPs had relatively small estimated effect sizes (S16 Fig), implying that there may be many more regions associated with asthma with increasingly smaller effect sizes that are missed by current GWAS sample sizes.

As a proxy for sensitivity, we calculated the proportion of FDR significant SNPs in the UK Biobank data set that were also found to be FDR significant both before (“iteration 0”) and after each iteration of Flexible cFDR. We found that the sensitivity increased from 0.127 to 0.131 after iteration 1 (leveraging average H3K27ac fold change values in lymphoid and CD56 cell types) and to 0.133 after iteration 2 (leveraging average H3K27ac fold change values in lung tissue and monocytes) (Fig 3E). As a proxy for specificity, we calculated the proportion of SNPs not FDR significant in the UK Biobank data set that were also not FDR significant both before (“iteration 0”) and after each iteration of Flexible cFDR, finding that the specificity remained close to 1 (≥ 0.9999975) (Fig 3F). We also found that the order of which we iterated over the auxiliary data had minimal impact on the results (S17 Fig).

At the locus level, 18 loci were FDR significant (*FDR* ≤ 0.000148249) in the original asthma GWAS ^42^. When used to leverage H3K27ac fold change values, Flexible cFDR identified 4 additional significant loci with index SNPs: rs9501077 (chr6:31167512), rs4148869 (chr6:32806576), rs9467715 (chr6:26341301) and rs167769 (chr12:57503775) (Fig ; Table 4). Three of the four (rs4148869, rs9467715 and rs167769) validated in the UK Biobank data set at Bonferroni corrected significance (for 4 tests the Bonferroni corrected significance threshold corresponding to α = 0.05 is 0.05*/*4 = 0.0125). One locus was found to be no longer FDR significant with index SNP rs12543811 (chr8:81278885).

**Table 4:** Summary of newly significant asthma index SNPs when using Flexible cFDR to leverage H3K27ac data. Details of index SNPs that became newly FDR significant (*FDR* < 0.000148249) after using Flexible cFDR to leverage H3K27ac fold change values with asthma GWAS *p*-values. Table contains the rsIDs (SNP), genomic positions (Chr: chromosome, BP: base pair), reference (Ref) and alternative (Alt) alleles, log ORs (beta), standard errors (SE) and *p*-values from the discovery GWAS, mean H3K27ac fold change values across asthma relevant cell types, *p*-values from UK Biobank and resultant *v*-values from Flexible cFDR. For the original *p*-values (and *v*-values), the corresponding FDR values are also given, calculated using the BH procedure.

| SNP | Chr | BP | Ref | Alt | beta | SE | H3K27ac percentile<sup>a</sup> | p | FDR (p) | p (UKBB) | v | FDR (v) |
| --- | --- | --- | --- | --- | --- | --- | --- | --- | --- | --- | --- | --- |
| rs167769 | 12 | 57503775 | C | T | 7.87e-02 | 1.50e-02 | 98.4th | 1.55e-07 | 4.04e-04 | 4.69e-24 | 3.75e-09 | 1.51e-05 |
| rs9467715 | 6 | 26341301 | T | C | -8.61e-02 | 1.61e-02 | 67.9th | 8.96e-08 | 2.49e-04 | 5.93e-04 | 3.83e-08 | 1.15e-04 |
| rs9501077 | 6 | 31167512 | A | G | -8.06e-02 | 1.54e-02 | 90.5th | 1.53e-07 | 3.99e-04 | 2.01e-02 | 1.91e-08 | 6.26e-05 |
| rs4148869 | 6 | 32806576 | C | T | 7.03e-02 | 1.39e-02 | 99.6th | 4.03e-07 | 9.28e-04 | 1.49e-15 | 9.07e-09 | 3.22e-05 |

<sup>a</sup> Mean H3K27ac percentile across cell types.

SNPs rs9501077 and rs4148869 reside in the major histocompatibility complex (MHC) region of the genome, which is renowned for its strong long-range LD structures that make it difficult to dissect genetic architecture in this region. rs9501077 and rs4148869 are in linkage equilibrium (*r*^2^ = 0.001), and are in very weak LD with the index SNP for the whole MHC region (rs9268969; *FDR* = 7.35*e* − 15; *r*^2^ = 0.005 and *r*^2^ = 0.001 respectively). rs9501077 (*p* = 1.53*e* − 07) has relatively high H3K27ac counts in asthma relevant cell types (mean percentile is 90th) and Flexible cFDR uses this extra disease-relevant information to increase the significance of this SNP beyond the significance threshold (FDR before Flexible cFDR = 3.99*e* − 04, FDR after Flexible cFDR = 6.26*e* − 05; Table 4). rs9501077 is found in the long non-coding RNA (lncRNA) gene, *HCG27* (HLA Complex Group 27), which has been linked to psoriasis ^73^, however the finding of a significant association with asthma for this SNP was not replicated in the UK Biobank data (UK Biobank *p* = 0.020).

SNP rs4148869 has very high H3K27ac fold change values in asthma relevant cell types (mean percentile is 99.6th) and so Flexible cFDR decreases the FDR value for this SNP from 9.28*e* – 04 to 3.22*e* − 05 when leveraging this auxiliary data (Table 4). This SNP is a 5’ UTR variant in the *TAP2* gene. The protein TAP2 assembles with TAP1 to form a transporter associated with antigen processing (TAP) complex. The TAP complex transports foreign peptides to the endoplasmic reticulum where they attach to MHC class I proteins which in turn are trafficked to the surface of the cell for antigen presentation to initiate an immune response ^74^. Studies have found *TAP2* to be associated with various immune-related disorders, including autoimmune thyroiditis and type 1 diabetes ^75,76^, and pulmonary tuberculosis in Iranian populations ^77^. Recently, Ma and colleagues ^78^ identified three cis-regulatory eSNPS for *TAP2* as candidates for childhood-onset asthma risk (rs9267798, rs4148882 and rs241456). One of these (rs4148882) is present in the asthma GWAS data set used for our analysis (*FDR* = 0.12) and is in weak LD with rs4148869 (*r*^2^ = 0.4).

SNP rs9467715 is a regulatory region variant with a raw FDR value that is very nearly significant in the original GWAS (FDR = 2.49*e* − 04 compared with FDR threshold of 1.48*e* − 04 used to call significant SNPs). This SNP has moderate H3K27ac fold change values in asthma relevant cell types (mean percentile is 67.9th) so that when these are leveraged using Flexible cFDR, the SNP is just pushed past the FDR significance threshold (FDR after Flexible cFDR = 1.15*e* − 04; Table 4).

SNP rs167769 has a borderline FDR value in the original GWAS discovery data set (*FDR* = 4.04*e* − 04) but was found to be significant in the multi-ancestry analysis in the same manuscript (*FDR* = 1.61*e* − 05) ^42^. This SNP has very high H3K27ac fold change values in asthma relevant cell types (mean percentile is 98.4th) and Flexible cFDR decreases the FDR value for this SNP to 1.51*e* − 05 when leveraging this auxiliary data (Table 4). rs167769 is an intron variant in *STAT6*, a gene that is activated by cytokines IL-4 and IL-13^79,80^ to initiate a Th2 response and ultimately inhibit transcribing of innate immune response genes ^81,82^. Transgenic mice over-expressing constitutively active *STAT6* in T cells are predisposed towards Th2 responses and allergic inflammation ^83,84^ whilst *STAT6*-knockout mice are protected from allergic pulmonary manifestations ^85^. Accordingly, rs167769 is strongly associated with *STAT6* expression in the blood ^86,87,88^ and lungs ^89^ and is associated with increased risk of childhood atopic dermatitis ^90,91^, which often progresses to allergic airways diseases such as asthma in adulthood. No genetic variants in the *STAT6* gene region (chr12:57489187-57525922) were identified as significant in the original GWAS, and only rs167769 was identified as significant after leveraging H3K27ac data using Flexible cFDR (S18 Fig).

One significant index SNP was no longer significant after applying Flexible cFDR. rs12543811 is located between genes *TPD52* and *ZBTB10* and has moderate H3K27ac fold change values in asthma relevant cell types (mean percentile is 52th). This SNP only just exceeds the FDR significance threshold in the original GWAS (FDR = 1.08*e* − 04 compared with FDR threshold of 1.48*e* − 04 used to call significant SNPs) but by leveraging its H3K27ac fold change values using Flexible cFDR, the resultant FDR value is just below the significance threshold (FDR after Flexible cFDR = 3.04*e* − 04; S19 Fig). This SNP is in strong LD with rs7009110 (*r*^2^ = 0.79) which has previous been associated with asthma plus hay fever but not with asthma alone ^92^. Conditional analyses show that these two SNPs represent the same signal which is likely to be associated with allergic asthma ^42^. rs12543811 was found to be significant in the UK Biobank data (UK Biobank *p* = 1.42*e* − 19).

### Comparison with existing methods

#### Boca and Leek’s FDR regression

We compared the results from Flexible cFDR when leveraging cell-type specific ChIP-seq data to those from BL when leveraging the exact same auxiliary data (Fig 5; S2 Table). The estimated probabilities that each SNP was null (not associated), which are calculated as an intermediate step in the method, ranged from 0.746 to 1 and were negatively correlated with H3K27ac fold change values in asthma relevant cell types (S20 Fig). In total, BL identified five SNPs as newly FDR significant, which replicate in the UK Biobank validation data set (rs4705950 UK Biobank *p* = 7.2*e* − 23, rs9268831 UK Biobank *p* = 1.3*e* − 42, rs17533090 UK Biobank *p* = 4.4*e* − 41, rs1871665 UK Biobank *p* = 5.5*e* − 24 and rs16924428 UK Biobank *p* = 3.6*e* − 40) (Fig 5B). These SNPs are a subset of the 69 newly significant SNPs identified by Flexible cFDR, except for rs16924428 which has very low H3K27ac fold change values in asthma relevant cell types (average value = 0.087). The sensitivity increased slightly from 0.127 to 0.128 after applying BL (compared to 0.133 after Flexible cFDR) (Fig 5C) and the specificity remained stable at 0.9999995 (Fig 5D). No SNPs were FDR significant in the discovery data set and no longer FDR significant after applying BL, and no new loci were found to be newly FDR significant (or newly not FDR significant).

**Figure 4:**
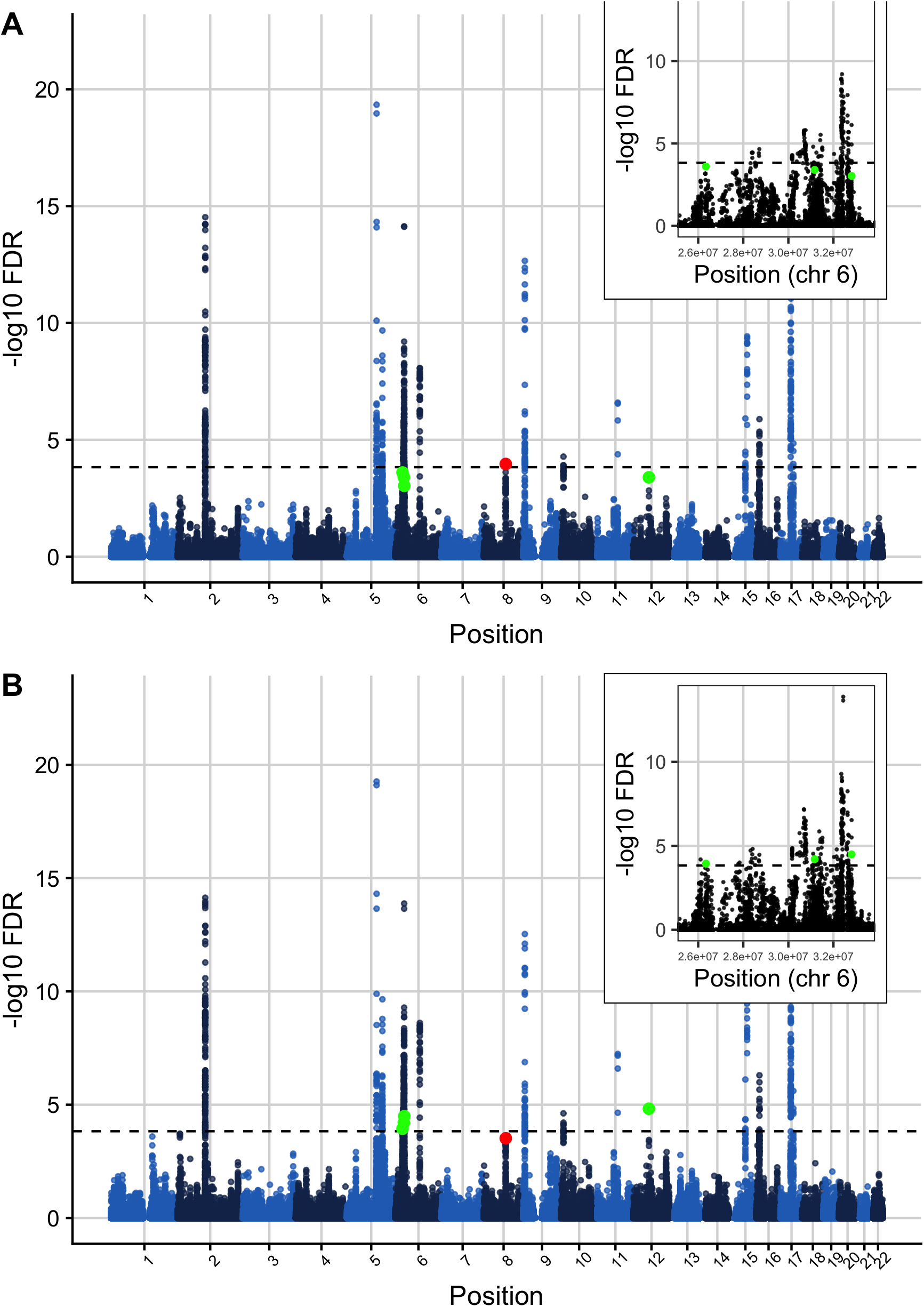
Manhattan plot of FDR values before and after applying Flexible cFDR to leverage H3K27ac data with asthma GWAS *p*-values. Manhattan plots of −log10 FDR values before (A) and after (B) applying Flexible cFDR leveraging H3K27ac counts in asthma relevant cell types. Points are coloured by chromosome and green points indicate the four index SNPs that are newly identified as FDR significant after Flexible cFDR (rs167769, rs9467715, rs9501077 and rs4148869) whilst the red point indicates the single index SNP that was newly identified as not FDR significant by Flexible cFDR (rs12543811). 30 Black dashed line at FDR significance threshold [−*log*10(0.000148249)].

**Figure 5:**
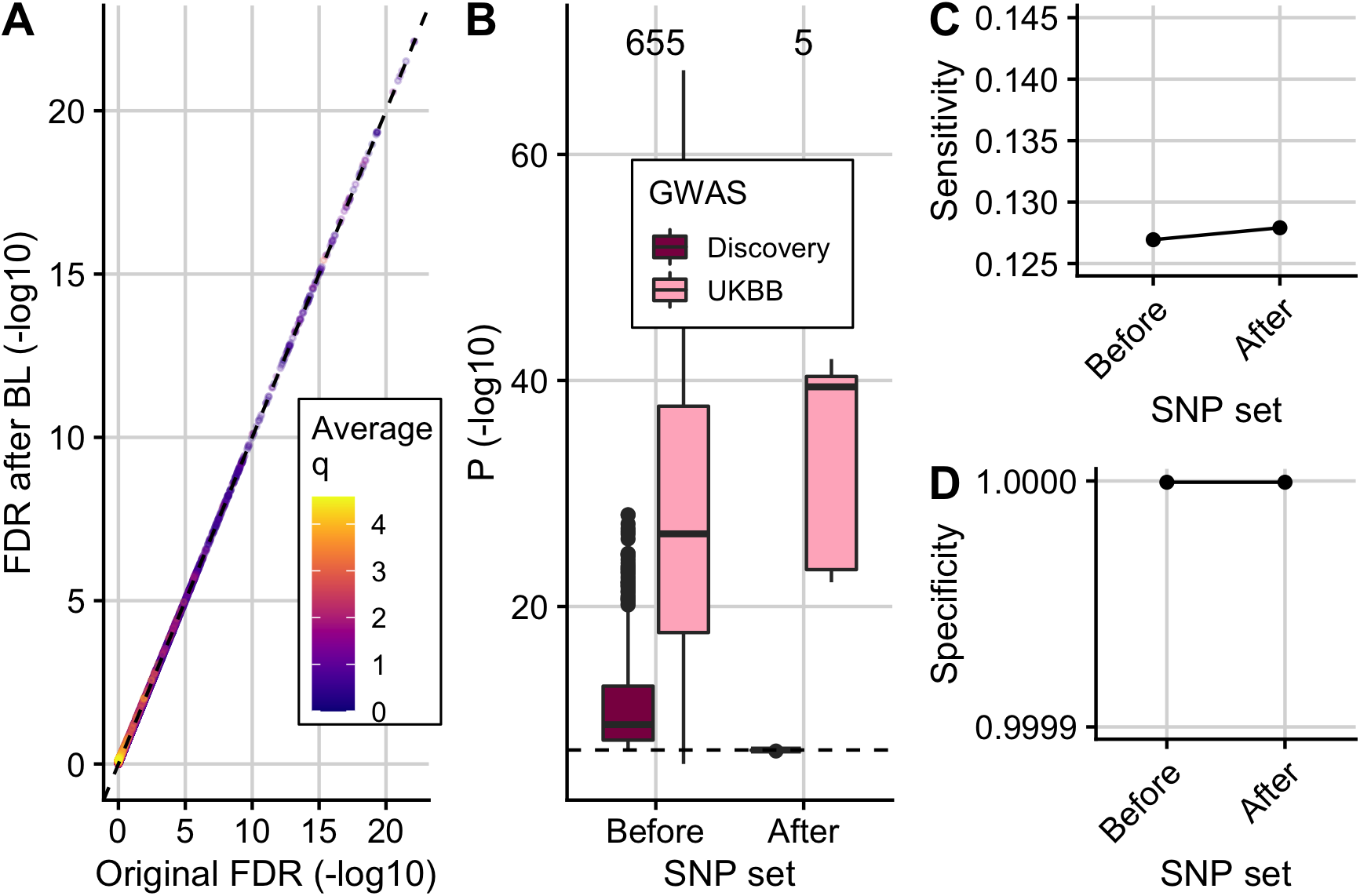
Using Boca and Leek’s FDR regression to leverage H3K27ac data with asthma GWAS *p*-values. (A) (−log10) adjusted *p*-values from BL against raw (-log10) FDR values coloured by average value of *q* (H3K27ac fold change value). (B) Box plots of (-log10) *p*-values in the discovery GWAS data set and the UK Biobank data set for the 655 SNPs that were FDR significant in the original GWAS (“before”) and 5 newly significant SNPs after applying BL (“after”). Black dashed line at genome-wide significance threshold (5*e* − 08). (C) Sensitivity and (D) specificity proxies for the results. Sensitivity proxy is calculated as the proportion of SNPs that are FDR significant in the UK Biobank data set that are also FDR significant in the original GWAS or after applying BL. Specificity is calculated as the proportion of SNPs that are not FDR significant in the UK Biobank data set that are also not FDR significant in the original GWAS or after BL.

#### FINDOR

We next compared results from Flexible cFDR when leveraging cell-type specific ChIP-seq data to those from FINDOR, which leverages a wider range of non-cell-type-specific functional annotations (S3 Table). FINDOR identified 119 newly FDR significant SNPs which had a median *p*-value of 4.44*e* − 15 in the UK Biobank validation data, but the maximum UK Biobank *p*-value for these 119 newly significant SNPs was 0.98 (Fig 6A; Fig 6B). The proportion of FDR significant SNPs in the UK Biobank data set that were also FDR significant in the discovery GWAS data set increased from 0.127 to 0.146 (compared to 0.128 after BL and 0.133 after Flexible cFDR) (Fig 6C) and the specificity remained ≥ 0.99999 (Fig 6D). The increase in sensitivity from FINDOR is greater than that of Flexible cFDR and BL, which may reflect the information gain in leveraging 96 annotations rather than a single histone mark.

**Figure 6:**
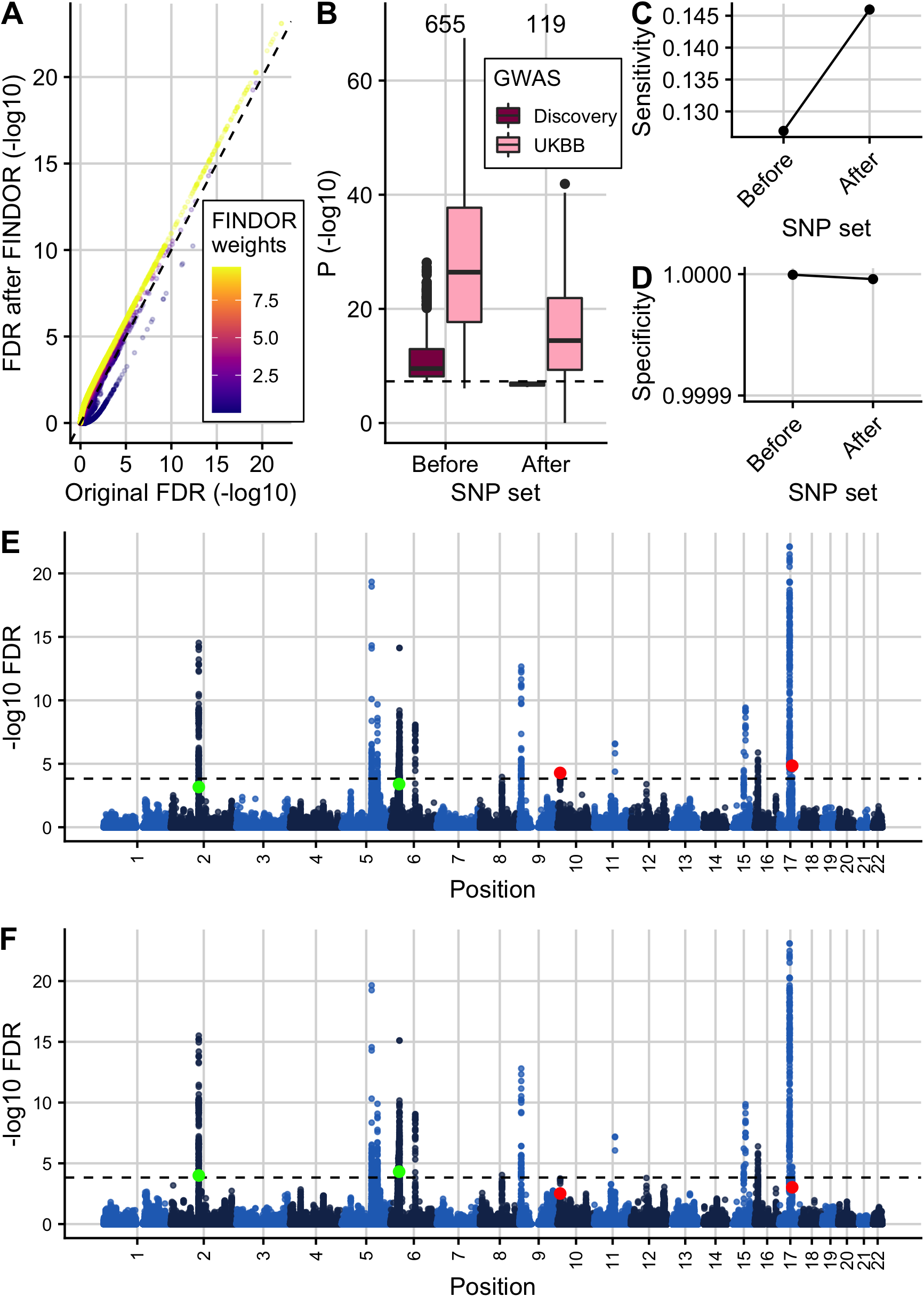
Results from FINDOR re-weighting of asthma GWAS *p*-values leveraging 96 baseline-LD model annotations. (A) (-log10) FDR values from FINDOR against (-log10) original FDR values coloured by FINDOR weights. (B) Box plots of (-log10) *p*-values in the discovery GWAS data set and the UK Biobank data set for the 655 SNPs that were FDR significant in the original GWAS (“before”) and 119 newly significant SNPs after re-weighting using FINDOR (“after”). Black dashed line at genome-wide significance threshold (5*e* − 08). (C) Sensitivity and specificity proxies for the FINDOR results. Sensitivity proxy is calculated as the proportion of SNPs that are FDR significant in the UK Biobank data set that are also FDR significant in the original GWAS or after *p*-value re-weighting using FINDOR. Specificity is calculated as the proportion of SNPs that are not FDR significant in the UK Biobank data set that are also not FDR significant in the original GWAS or after *p*-value re-weighting using FINDOR. Manhattan plots of FDR values before (E) and after (F) re-weighting by FINDOR. Green points indicate the two index SNPs that are newly identified as FDR significant by FINDOR [rs13018263 (chr2:103092270) and rs9501077 (chr6:31167512)]. Red points indicate the two index SNPs that are newly identified as not FDR significant by FINDOR [rs2589561 (chr10:9046645) and rs17637472 (chr17:47461433)]. Black dashed line at FDR significance threshold [−*log*10(0.000148249)].

At the locus level, FINDOR identified two newly FDR significant index SNPs: rs13018263 (chr2:103092270; original *FDR* = 6.79*e*−04, new *FDR* = 1.00*e*−04) and rs9501077 (chr6:31167512; original *FDR* = 3.99*e* − 04, new *FDR* = 4.86*e* − 05) (Fig 6E; Fig 6F). SNP rs13018263 is an intronic variant in *SLC9A4* and is strongly significant in the UK Biobank validation data set (*p* = 4.78*e* − 31). Ferreira and colleagues ^93^ highlighted rs13018263 as a potential eQTL for *IL18RAP*, a gene which is involved in IL-18 signalling which in turn mediates Th1 responses ^94^, and is situated just upstream of *SLC9A4*. Genetic variants in *IL18RAP* are associated with many immune-mediated diseases, including atopic dermatitis^95^ and type 1 diabetes ^96^. Interestingly, although different auxiliary data was leveraged using Flexible cFDR and FINDOR in our analyses, both methods found index SNP rs9501077 to be newly significant, but this SNP did not validate in the UK Biobank data (UK Biobank *p* = 0.020).

Two additional index SNPs were found to be no longer significant after re-weighting by FINDOR, rs2589561 (chr10:9046645; original *FDR* = 5.25*e* − 05, new *FDR* = 3.06*e* − 03) and rs17637472 (chr17:47461433; original *FDR* = 1.42*e* −05, new *FDR* = 9.42*e* −04), however both of these SNPs were strongly significant in the UK Biobank validation data set (*p* = 2.09*e* − 29 and *p* = 1.75*e* − 14 respectively).

SNP rs2589561 is a gene desert that is 929kb from *GATA3*, a transcription factor of the Th2 pathway which mediates the immune response to allergens ^42,97^. Hi-C data in hematopoietic cells showed that two proxies of rs2589561 (*r*^2^ > 0.9) are located in a region that interacts with the *GATA3* promoter in CD4+ T cells ^98^, suggesting that rs2589561 could function as a distal regulator of *GATA3* in this asthma relevant cell type. rs2589561 has relatively high H3K27ac fold change values in the asthma relevant cell types leveraged by Flexible cFDR (mean percentile is 85th) and Flexible cFDR decreased the FDR value from 5.25*e* − 05 to 2.41*e* − 05.

SNP rs17637472 is a strong cis-eQTL for *GNGT2* in blood ^86,87,88,99^, a gene whose protein product is involved in NF-*κ*B activation ^100^. This SNP has moderate H3K27ac fold change values in relevant cell types (mean percentile is 62th) and the FDR values for this SNP were similar both before and after using Flexible cFDR to leverage the H3K27ac data (original *FDR* = 1.42*e* − 05, new *FDR* = 1.46*e* − 05).

## Discussion

Developments in molecular biology have enabled researchers to decipher the functional effects of various genomic signatures. We are now in a position to prioritise sequence variants associated with various phenotypes not just by their genetic association statistics but also based on our biological understanding of their functional role. Originally designed for the specific purpose of leveraging test statistics from genetically related traits, we have extended the cFDR framework ^41^ to support auxiliary data from arbitrary continuous distributions. Our extension, Flexible cFDR, provides a statistically robust framework to leverage functional genomic data with genetic association statistics to boost power for GWAS discovery.

We compared the performance of Flexible cFDR to that of four comparator methods which have previously been shown to outperform other approaches ^43,15^: GenoWAP, IHW, BL and FINDOR. We also tried to compare our method to AdaPT ^10^, but this approach uses a *p*-value masking procedure which takes many iterations of optimisation and can be computationally expensive ^101^. We found AdaPT to be too computationally demanding for large-scale GWAS data and previous studies suggest that a SNP pre-filtering stage is required ^36^. Of the methods considered, we found that only BL was as versatile as Flexible cFDR. Specifically, IHW currently only supports univariate covariates and, unlike Flexible cFDR, cannot be applied iteratively to leverage multi-dimensional covariates. In GenoWAP, the prior probabilities used in the model are calculated as the mean GenoCanyon score (or tissue-specific GenoSkyline ^102^ or GenoSkyline-Plus ^103^ score) of the surrounding 10, 000 base pairs, thereby restricting its utility to leveraging only these scores (which we found were unlikely to capture enough disease relevant information to substantially alter conclusions from a study). Whilst in FINDOR, SNPs are binned based on how well they tag heritability enriched categories and this requires the estimation of *χ*^2^ statistics (i.e., tagged variance) for each SNP using a range of functional annotations, which are generally those in the baselineLD model ^27^. Users are thus required to run LD-score regression prior to running FINDOR, and this two-step approach may limit the accessibility of the method. Although BL was as versatile as Flexible cFDR, we found that it often failed to control the FDR and was less powerful than Flexible cFDR in a simulation-based analysis. Whilst FINDOR was shown to be the most powerful method, this may reflect the information gain in leveraging 96 annotations rather than a single histone mark. This emphasises the importance of being able to iterate over different auxiliary measures, and suggests that a fruitful area of extension for cFDR will be to increase the robustness of FDR control for dependent *q* across multiple iterations.

Flexible cFDR has several key advantages over competing methods. It does not bin variables and does not rely on subjective thresholding or normalised weighting schemes, which hinder many of the existing methods ^9,4,25,5,6,15,3,14^. Moreover, Flexible cFDR outputs quantities analogous to *p*-values which can be used directly in any error-rate controlling procedure, and which also permit iteration to support multi-dimensional covariate data. Whilst LD between SNPs is often a concern (e.g. because methods such as KDE assume independence between observations), we fit the KDE to a subset of LD-independent SNPs but then generate *v*-values for the full set of SNPs, thereby benefiting from computational efficiency but also facilitating downstream analyses which typically require the full set of SNPs, such as fine-mapping or meta-analysis. LD means that the *v*-values will be positively correlated, so we appeal to the established robustness of the Benjamini-Hochberg FDR estimation to positive dependency ^104^.

Whilst larger case and control cohort sizes will boost statistical power for GWAS discovery, incorporating functional data provides an additional layer of biological evidence that an increase in sample sizes alone cannot provide. There are also instances in the rare disease domain where case sample sizes are restricted by the number of cases available for recruitment (for example in primary immunodeficiency disorder ^105^), and our method has potential utility in these instances as it provides an alternative approach to increase statistical power. The choice of functional data to use may be guided by prior knowledge, or in a data driven manner using a method such as GARFIELD ^106^ to quantify the enrichment of GWAS signals in different functional marks. Moreover, our method intrinsically evaluates the relevance of the auxiliary data by comparing the joint probability density of the test statistics and the auxiliary data to the joint density assuming independence, and can therefore be used to inform researchers of relevant functional signatures and cell types.

Our manuscript describes four key advances enabling the extension of the cFDR framework to the functional genomics setting. Firstly, we derive an estimator based on a 2-dimensional KDE of the bivariate distribution rather than empirical estimates, making our method considerably faster than earlier empirical approaches. Secondly, the cFDR framework requires the estimation of 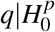, which Liley and Wallace (2021) ^41^ approximate by *q*|*P* > 1/2. In contrast, Flexible cFDR utilises the local FDR to empirically evaluate the influence of specific *p*-value quantities on the null hypothesis and uses these in the estimation of 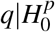. Thirdly, we remove the assumption that 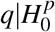 can be transformed to a mixture of centred normals, and instead integrate over the previously estimated KDE, which relaxes the distributional assumptions placed on the auxiliary data. Finally, Flexible cFDR is supported by user-oriented software documented on an easy-to-navigate website (https://annahutch.github.io/fcfdr/). The website features several fully reproducible vignettes which illustrate how the method can be applied to a particular data set at the desired level of error control. We hope this support will make Flexible cFDR accessible to a wider range of researchers.

One can see that the scale on which the auxiliary data is measured may impact the performance of Flexible cFDR. Usual concerns about KDE apply, including that data with hard boundaries should be transformed [e.g. the logit function can transform data on (0, 1) to (−∞, +∞)]. Further, it is likely to be important that the auxiliary data is on a scale such that the changing relationship with *p* can be adequately captured with a reasonable bandwidth. For example, if the auxiliary data are GWAS *p*-values, we would recommend conditioning not on the raw *p*-values, but on log transformed values. The optimal scale for the auxiliary data is likely to depend on the relationship between the principal *p*-values and the auxiliary data, and is not something we have explored here, but as usual, data visualisation is likely to be helpful to confirm that the scale for the auxiliary data is sensible.

Our method has several limitations. Firstly, care must be taken to ensure that the auxiliary data to be leveraged iteratively is capturing distinct disease-relevant features to prevent multiple adjustment using the same auxiliary data. The definition of “distinct disease-relevant features” to leverage is at the user’s discretion and sparks an interesting philosophical discussion. For example, leveraging data iteratively from various genomic assays measuring the same genomic feature at different resolutions may be deemed invalid for some researchers but valid for others, since if the mark is repeatedly identified by different assays then it is more likely to be reliably present. Whilst we show that our method is robust to minor departures from 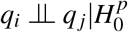, this does not extend to strongly related *q*. We would argue that the conservative approach would be to average over correlated auxiliary data, to ensure that the *q* vectors are not strongly correlated.

Secondly, the cFDR framework assumes a positive stochastically monotonic relationship between the test statistics and the auxiliary data: specifically, low *p*-values are enriched for low values in the auxiliary data. Our method automatically calculates the correlation between *p* and *q* and if this is negative then the auxiliary data is transformed to *q* := −*q*. However, if the relationship is non-monotonic (for example low *p*-values are enriched for both very low and very high values in the auxiliary data) then the cFDR framework cannot simultaneously shrink *v*-values for these two extremes. This non-monotonic relationship is unlikely when leveraging single functional genomic marks, but may occur if, for example, multiple marks were decomposed via PCA. We therefore recommend that users use the `corr_plot` and `stratified_qqplot` functions in the `fcfdr` R package to visualise the relationship between the relationship between the two data types. Note that this restriction could be removed if we used density instead of distribution functions, and worked at the level of local FDR ^45^, but this would in turn reduce the robustness our method has to data sparsity in the (*p, q*) plane.

Finally, in our asthma application we only leveraged data for a single histone modification across various cell types. Additional data measuring other histone modifications (e.g. repressive marks) could also be leveraged to further increase power.

Overall, we anticipate that Flexible cFDR will be a valuable tool to leverage functional genomic data with GWAS test statistics to boost power for GWAS discovery.

## Web Resources

Code to reproduce results from this manuscript https://github.com/annahutch/fcfdr_manuscript

Flexible cFDR software https://github.com/annahutch/fcfdr

Empirical cFDR software https://github.com/jamesliley/cfdr/

GWAS Catalog https://www.ebi.ac.uk/gwas/

Asthma GWAS summary statistics ftp://ftp.ebi.ac.uk/pub/databases/gwas/summary_statistics/DemenaisF_29273806_GCST006862/harmonised/29273806-GCST006862-EFO_0000270.h.tsv.gz

Neale lab UK Biobank http://www.nealelab.is/uk-biobank

Neale lab asthma summary statistics https://www.dropbox.com/s/kp9bollwekaco0s/20002_1111.gwas.imputed_v3.both_sexes.tsv.bgz?dl=0

NIH Roadmap data resource https://www.ncbi.nlm.nih.gov/geo/roadmap/epigenomics/?search=pbmc&display=200

NIH Roadmap H3K27ac data resource https://egg2.wustl.edu/roadmap/data/byFileType/signal/consolidated/macs2signal/foldChange/

FINDOR software https://github.com/gkichaev

GenoCanyon annotations http://genocanyon.med.yale.edu/GenoCanyon_Downloads.html

GenoWAP software https://github.com/rlpowles/GenoWAP-V1.2

## References

[1] Sivakumaran S, Agakov F, Theodoratou E, Prendergast JG, Zgaga L, Manolio T, et al. Abundant Pleiotropy in Human Complex Diseases and Traits. American Journal of Human Genetics. 2011;89(5):607–618. doi:10.1016/j.ajhg.2011.10.004.

[2] Schork AJ, Thompson WK, Pham P, Torkamani A, Roddey JC, Sullivan PF, et al. All SNPs Are Not Created Equal: Genome-Wide Association Studies Reveal a Consistent Pattern of Enrichment among Functionally Annotated SNPs. PLoS genetics. 2013;9(4):e1003449. doi:10.1371/journal.pgen.1003449.

[3] Genovese CR, Roeder K, Wasserman L. False Discovery Control with P-Value Weighting. Biometrika. 2006;93(3):509–524. doi:10.1093/biomet/93.3.509.

[4] Sun L, Craiu RV, Paterson AD, Bull SB. Stratified False Discovery Control for Large-Scale Hypothesis Testing with Application to Genome-Wide Association Studies. Genetic Epidemiology. 2006;30(6):519–530. doi:10.1002/gepi.20164.

[5] Hu JX, Zhao H, Zhou HH. False Discovery Rate Control With Groups. Journal of the American Statistical Association. 2010;105(491):1215–1227. doi:10.1198/jasa.2010.tm09329.

[6] Ignatiadis N, Klaus B, Zaugg JB, Huber W. Data-Driven Hypothesis Weighting Increases Detection Power in Genome-Scale Multiple Testing. Nature Methods. 2016;13(7):577–580. doi:10.1038/nmeth.3885.

[7] Ferkingstad E, Frigessi A, Rue H, Thorleifsson G, Kong A. Unsupervised Empirical Bayesian Multiple Testing with External Covariates. The Annals of Applied Statistics. 2008;2(2):714–735. doi:10.1214/08-AOAS158.

[8] Basu P, Cai TT, Das K, Sun W. Weighted False Discovery Rate Control in Large-Scale Multiple Testing. Journal of the American Statistical Association. 2018;113(523):1172–1183. doi:10.1080/01621459.2017.1336443.

[9] Bourgon R, Gentleman R, Huber W. Independent Filtering Increases Detection Power for High-Throughput Experiments. Proceedings of the National Academy of Sciences. 2010;107(21):9546–9551. doi:10.1073/pnas.0914005107.

[10] Lei L, Fithian W. AdaPT: An Interactive Procedure for Multiple Testing with Side Information. arXiv:160906035 [stat]. 2018;.

[11] Li A, Barber RF. Multiple Testing with the Structure Adaptive Benjamini-Hochberg Algorithm. arXiv:160607926 [stat]. 2017;.

[12] Roeder K, Wasserman L. Genome-Wide Significance Levels and Weighted Hypothesis Testing. Statistical Science. 2009;24(4):398–413. doi:10.1214/09-STS289.

[13] Cai TT, Sun W, Wang W. Covariate-Assisted Ranking and Screening for Large-Scale Two-Sample Inference. Journal of the Royal Statistical Society: Series B (Statistical Methodology). 2019;81(2):187–234. doi:10.1111/rssb.12304.

[14] Lu Q, Yao X, Hu Y, Zhao H. GenoWAP: GWAS Signal Prioritization through Integrated Analysis of Genomic Functional Annotation. Bioinformatics. 2016;32(4):542–548. doi:10.1093/bioinformatics/btv610.

[15] Kichaev G, Bhatia G, Loh PR, Gazal S, Burch K, Freund MK, et al. Leveraging Polygenic Functional Enrichment to Improve GWAS Power. American Journal of Human Genetics. 2019;104(1):65–75. doi:10.1016/j.ajhg.2018.11.008.

[16] Pickrell JK. Joint Analysis of Functional Genomic Data and Genome-Wide Association Studies of 18 Human Traits. American Journal of Human Genetics. 2014;94(4):559–573. doi:10.1016/j.ajhg.2014.03.004.

[17] Hou L, Zhao H. A Review of Post-GWAS Prioritization Approaches. Frontiers in Genetics. 2013;4. doi:10.3389/fgene.2013.00280.

[18] Sveinbjornsson G, Albrechtsen A, Zink F, Gudjonsson SA, Oddson A, Másson G, et al. Weighting Sequence Variants Based on Their Annotation Increases Power of Whole-Genome Association Studies. Nature Genetics. 2016;48(3):314–317. doi:10.1038/ng.3507.

[19] Roeder K, Devlin B, Wasserman L. Improving Power in Genome-Wide Association Studies: Weights Tip the Scale. Genetic Epidemiology. 2007;31(7):741–747. doi:10.1002/gepi.20237.

[20] Eskin E. Increasing Power in Association Studies by Using Linkage Disequilibrium Structure and Molecular Function as Prior Information. Genome Research. 2008;18(4):653–660. doi:10.1101/gr.072785.107.

[21] Darnell G, Duong D, Han B, Eskin E. Incorporating Prior Information into Association Studies. Bioinformatics. 2012;28(12):i147–i153. doi:10.1093/bioinformatics/bts235.

[22] Wen X, Lee Y, Luca F, Pique-Regi R. Efficient Integrative Multi-SNP Association Analysis via Deterministic Approximation of Posteriors. American Journal of Human Genetics. 2016;98(6):1114–1129. doi:10.1016/j.ajhg.2016.03.029.

[23] Benjamini Y, Hochberg Y. Controlling the False Discovery Rate: A Practical and Powerful Approach to Multiple Testing. Journal of the Royal Statistical Society Series B (Methodological). 1995;57(1):289–300.

[24] Holm S. A Simple Sequentially Rejective Multiple Test Procedure. Scandinavian Journal of Statistics. 1979;6(2):65–70.

[25] Benjamini Y, Krieger AM, Yekutieli D. Adaptive Linear Step-up Procedures That Control the False Discovery Rate. Biometrika. 2006;93(3):491–507.

[26] Finucane HK, Bulik-Sullivan B, Gusev A, Trynka G, Reshef Y, Loh PR, et al. Partitioning Heritability by Functional Annotation Using Genome-Wide Association Summary Statistics. Nature Genetics. 2015;47(11):1228–1235. doi:10.1038/ng.3404.

[27] Gazal S, Finucane HK, Furlotte NA, Loh PR, Palamara PF, Liu X, et al. Linkage Disequilibrium–Dependent Architecture of Human Complex Traits Shows Action of Negative Selection. Nature Genetics. 2017;49(10):1421–1427. doi:10.1038/ng.3954.

[28] Storey JD, Tibshirani R. Statistical Significance for Genomewide Studies. Proceedings of the National Academy of Sciences. 2003;100(16):9440–9445. doi:10.1073/pnas.1530509100.

[29] Storey JD. A Direct Approach to False Discovery Rates. Journal of the Royal Statistical Society: Series B (Statistical Methodology). 2002;64(3):479–498. doi:10.1111/1467-9868.00346.

[30] Boca SM, Leek JT. A Direct Approach to Estimating False Discovery Rates Conditional on Covariates. PeerJ. 2018;6:e6035. doi:10.7717/peerj.6035.

[31] Efron B, Tibshirani R, Storey JD, Tusher V. Empirical Bayes Analysis of a Microarray Experiment. Journal of the American Statistical Association. 2001;96(456):1151–1160. doi:10.1198/016214501753382129.

[32] Cai TT, Sun W. Simultaneous Testing of Grouped Hypotheses: Finding Needles in Multiple Haystacks. Journal of the American Statistical Association. 2009;104(488):1467–1481. doi:10.1198/jasa.2009.tm08415.

[33] Ploner A, Calza S, Gusnanto A, Pawitan Y. Multidimensional Local False Discovery Rate for Microarray Studies. Bioinformatics (Oxford, England). 2006;22(5):556–565. doi:10.1093/bioinformatics/btk013.

[34] Barber RF, Candès EJ. Controlling the False Discovery Rate via Knockoffs. Annals of Statistics. 2015;43(5):2055–2085. doi:10.1214/15-AOS1337.

[35] Lei L, Fithian W. Power of Ordered Hypothesis Testing. arXiv:160601969 [stat]. 2016;.

[36] Yurko R, G’Sell M, Roeder K, Devlin B. A Selective Inference Approach for False Discovery Rate Control Using Multiomics Covariates Yields Insights into Disease Risk. Proceedings of the National Academy of Sciences. 2020;117(26):15028–15035. doi:10.1073/pnas.1918862117.

[37] Ignatiadis N, Huber W. Covariate Powered Cross-Weighted Multiple Testing. arXiv:170105179 [stat]. 2020;.

[38] Andreassen OA, Thompson WK, Schork AJ, Ripke S, Mattingsdal M, Kelsoe JR, et al. Improved Detection of Common Variants Associated with Schizophrenia and Bipolar Disorder Using Pleiotropy-Informed Conditional False Discovery Rate. PLOS Genetics. 2013;9(4):e1003455. doi:10.1371/journal.pgen.1003455.

[39] Andreassen OA, McEvoy Linda K, Thompson Wesley K, Wang Yunpeng, Reppe Sjur, Schork Andrew J, et al. Identifying Common Genetic Variants in Blood Pressure Due to Polygenic Pleiotropy With Associated Phenotypes. Hypertension. 2014;63(4):819–826. doi:10.1161/HYPERTENSIONAHA.113.02077.

[40] Andreassen OA, Harbo HF, Wang Y, Thompson WK, Schork AJ, Mattingsdal M, et al. Genetic Pleiotropy between Multiple Sclerosis and Schizophrenia but Not Bipolar Disorder: Differential Involvement of Immune-Related Gene Loci. Molecular Psychiatry. 2015;20(2):207–214. doi:10.1038/mp.2013.195.

[41] Liley J, Wallace C. Accurate Error Control in High-Dimensional Association Testing Using Conditional False Discovery Rates. Biometrical Journal. 2021; doi:10.1002/bimj.201900254.

[42] Demenais F, Margaritte-Jeannin P, Barnes KC, Cookson WOC, Altmüller J, Ang W, et al. Multiancestry Association Study Identifies New Asthma Risk Loci That Colocalize with Immune-Cell Enhancer Marks. Nature Genetics. 2018;50(1):42–53. doi:10.1038/s41588-017-0014-7.

[43] Korthauer K, Kimes PK, Duvallet C, Reyes A, Subramanian A, Teng M, et al. A Practical Guide to Methods Controlling False Discoveries in Computational Biology. Genome Biology. 2019;20(1):118. doi:10.1186/s13059-019-1716-1.

[44] Sudlow C, Gallacher J, Allen N, Beral V, Burton P, Danesh J, et al. UK Biobank: An Open Access Resource for Identifying the Causes of a Wide Range of Complex Diseases of Middle and Old Age. PLoS Medicine. 2015;12(3). doi:10.1371/journal.pmed.1001779.

[45] Efron B. Size, Power and False Discovery Rates. Annals of Statistics. 2007;35(4):1351–1377. doi:10.1214/009053606000001460.

[46] Wen X. A Unified View of False Discovery Rate Control: Reconciliation of Bayesian and Frequentist Approaches. arXiv:180305284 [stat]. 2018;.

[47] Liley J, Wallace C. A Pleiotropy-Informed Bayesian False Discovery Rate Adapted to a Shared Control Design Finds New Disease Associations From GWAS Summary Statistics. PLOS Genetics. 2015;11(2):e1004926. doi:10.1371/journal.pgen.1004926.

[48] Venables WN, Ripley BD. Modern Applied Statistics with S. 4th ed. Statistics and Computing. New York: Springer-Verlag; 2002.

[49] Speed D, Holmes J, Balding DJ. Evaluating and Improving Heritability Models Using Summary Statistics. Nature Genetics. 2020;52(4):458–462. doi:10.1038/s41588-020-0600-y.

[50] Chang CC, Chow CC, Tellier LC, Vattikuti S, Purcell SM, Lee JJ. Second-Generation PLINK: Rising to the Challenge of Larger and Richer Datasets. GigaScience. 2015;4(1). doi:10.1186/s13742-015-0047-8.

[51] Walter K, Min JL, Huang J, Crooks L, Memari Y, McCarthy S, et al. The UK10K Project Identifies Rare Variants in Health and Disease. Nature. 2015;526(7571):82–90. doi:10.1038/nature14962.

[52] Berisa T, Pickrell JK. Approximately Independent Linkage Disequilibrium Blocks in Human Populations. Bioinformatics. 2016;32(2):283–285. doi:10.1093/bioinformatics/btv546.

[53] Wellcome Trust Case Control Consortium. Genome-Wide Association Study of 14,000 Cases of Seven Common Diseases and 3,000 Shared Controls. Nature. 2007;447(7145):661–678. doi:10.1038/nature05911.

[54] Fortune MD, Wallace C. simGWAS: A Fast Method for Simulation of Large Scale Case–Control GWAS Summary Statistics. Bioinformatics. 2019;35(11):1901–1906. doi:10.1093/bioinformatics/bty898.

[55] Speed D, Cai N, Johnson MR, Nejentsev S, Balding DJ. Re-Evaluation of SNP Heritability in Complex Human Traits. Nature genetics. 2017;49(7):986–992. doi:10.1038/ng.3865.

[56] Leek JT, Jager L, Boca SM, Konopka T. Swfdr: Science-Wise False Discovery Rate and Proportion of True Null Hypotheses Estimation; 2021. Bioconductor version: Release (3.12).

[57] Buniello A, MacArthur JAL, Cerezo M, Harris LW, Hayhurst J, Malangone C, et al. The NHGRI-EBI GWAS Catalog of Published Genome-Wide Association Studies, Targeted Arrays and Summary Statistics. Nucleic Acids Research. 2019;47(D1):D1005–D1012. doi:10.1093/nar/gky1120.

[58] Kuhn RM, Haussler D, Kent WJ. The UCSC Genome Browser and Associated Tools. Briefings in Bioinformatics. 2013;14(2):144. doi:10.1093/bib/bbs038.

[59] Auton A, Abecasis GR, Altshuler DM, Durbin RM, Abecasis GR, Bentley DR, et al. A Global Reference for Human Genetic Variation. Nature. 2015;526(7571):68–74. doi:10.1038/nature15393.

[60] Boyle AP, Hong EL, Hariharan M, Cheng Y, Schaub MA, Kasowski M, et al. Annotation of Functional Variation in Personal Genomes Using RegulomeDB. Genome Research. 2012;22(9):1790–1797. doi:10.1101/gr.137323.112.

[61] Khurana E, Fu Y, Colonna V, Mu XJ, Kang HM, Lappalainen T, et al. Integrative Annotation of Variants from 1092 Humans: Application to Cancer Genomics. Science. 2013;342(6154). doi:10.1126/science.1235587.

[62] Ritchie GRS, Dunham I, Zeggini E, Flicek P. Functional Annotation of Noncoding Sequence Variants. Nature Methods. 2014;11(3):294–296. doi:10.1038/nmeth.2832.

[63] Kircher M, Witten DM, Jain P, O’Roak BJ, Cooper GM, Shendure J. A General Framework for Estimating the Relative Pathogenicity of Human Genetic Variants. Nature Genetics. 2014;46(3):310–315. doi:10.1038/ng.2892.

[64] Lu Q, Hu Y, Sun J, Cheng Y, Cheung KH, Zhao H. A Statistical Framework to Predict Functional Non-Coding Regions in the Human Genome Through Integrated Analysis of Annotation Data. Scientific Reports. 2015;5(1):10576. doi:10.1038/srep10576.

[65] Creyghton MP, Cheng AW, Welstead GG, Kooistra T, Carey BW, Steine EJ, et al. Histone H3K27ac Separates Active from Poised Enhancers and Predicts Developmental State. Proceedings of the National Academy of Sciences of the United States of America. 2010;107(50):21931. doi:10.1073/pnas.1016071107.

[66] Corradin O, Scacheri PC. Enhancer Variants: Evaluating Functions in Common Disease. Genome Medicine. 2014;6(10). doi:10.1186/s13073-014-0085-3.

[67] Bernstein BE, Stamatoyannopoulos JA, Costello JF, Ren B, Milosavljevic A, Meissner A, et al. The NIH Roadmap Epigenomics Mapping Consortium. Nature biotechnology. 2010;28(10):1045–1048. doi:10.1038/nbt1010-1045.

[68] Quinlan AR, Hall IM. BEDTools: A Flexible Suite of Utilities for Comparing Genomic Features. Bioinformatics. 2010;26(6):841–842. doi:10.1093/bioinformatics/btq033.

[69] Villar D, Berthelot C, Aldridge S, Rayner TF, Lukk M, Pignatelli M, et al. Enhancer Evolution across 20 Mammalian Species. Cell. 2015;160(3):554–566. doi:10.1016/j.cell.2015.01.006.

[70] Marnetto D, Mantica F, Molineris I, Grassi E, Pesando I, Provero P. Evolutionary Rewiring of Human Regulatory Networks by Waves of Genome Expansion. The American Journal of Human Genetics. 2018;102(2):207–218. doi:10.1016/j.ajhg.2017.12.014.

[71] Hujoel MLA, Gazal S, Hormozdiari F, van de Geijn B, Price AL. Disease Heritability Enrichment of Regulatory Elements Is Concentrated in Elements with Ancient Sequence Age and Conserved Function across Species. The American Journal of Human Genetics. 2019;104(4):611–624. doi:10.1016/j.ajhg.2019.02.008.

[72] Soskic B, Cano-Gamez E, Smyth DJ, Rowan WC, Nakic N, Esparza-Gordillo J, et al. Chromatin Activity at GWAS Loci Identifies T Cell States Driving Complex Immune Diseases. Nature Genetics. 2019;51(10):1486–1493. doi:10.1038/s41588-019-0493-9.

[73] Villarreal-Martínez A, Gallardo-Blanco H, Cerda-Flores R, Torres-Muñoz I, Gómez-Flores M, Salas-Alanís J, et al. Candidate Gene Polymorphisms and Risk of Psoriasis: A Pilot Study. Experimental and Therapeutic Medicine. 2016;11(4):1217–1222. doi:10.3892/etm.2016.3066.

[74] Hewitt EW. The MHC Class I Antigen Presentation Pathway: Strategies for Viral Immune Evasion. Immunology. 2003;110(2):163–169. doi:10.1046/j.1365-2567.2003.01738.x.

[75] Tomer Y, Dolan LM, Kahaly G, Divers J, D’Agostino RB, Imperatore G, et al. Genome Wide Identification of New Genes and Pathways in Patients with Both Autoimmune Thyroiditis and Type 1 Diabetes. Journal of Autoimmunity. 2015;60:32–39. doi:10.1016/j.jaut.2015.03.006.

[76] Carvalho-Silva D, Pierleoni A, Pignatelli M, Ong C, Fumis L, Karamanis N, et al. Open Targets Platform: New Developments and Updates Two Years On. Nucleic Acids Research. 2019;47(D1):D1056–D1065. doi:10.1093/nar/gky1133.

[77] Naderi M, Hashemi M, Amininia S. Association of TAP1 and TAP2 Gene Polymorphisms with Susceptibility to Pulmonary Tuberculosis. Iranian Journal of Allergy, Asthma and Immunology. 2016; p. 62–68.

[78] Ma X, Wang P, Xu G, Yu F, Ma Y. Integrative Genomics Analysis of Various Omics Data and Networks Identify Risk Genes and Variants Vulnerable to Childhood-Onset Asthma. BMC Medical Genomics. 2020;13(1):123. doi:10.1186/s12920-020-00768-z.

[79] Takeda K, Kamanaka M, Tanaka T, Kishimoto T, Akira S. Impaired IL-13-Mediated Functions of Macrophages in STAT6-Deficient Mice. The Journal of Immunology. 1996;157(8):3220–3222.

[80] Takeda K, Tanaka T, Shi W, Matsumoto M, Minami M, Kashiwamura S, et al. Essential Role of Stat6 in IL-4 Signalling. Nature. 1996;380(6575):627–630. doi:10.1038/380627a0.

[81] Ohmori Y, Hamilton TA. Interleukin-4/STAT6 Represses STAT1 and NF-Kappa B-Dependent Transcription through Distinct Mechanisms. The Journal of Biological Chemistry. 2000;275(48):38095–38103. doi:10.1074/jbc.M006227200.

[82] Albanesi C, Fairchild HR, Madonna S, Scarponi C, Pità OD, Leung DYM, et al. IL-4 and IL-13 Negatively Regulate TNF-*α*- and IFN-*γ*-Induced *β*-Defensin Expression through STAT-6, Suppressor of Cytokine Signaling (SOCS)-1, and SOCS-3. The Journal of Immunology. 2007;179(2):984–992. doi:10.4049/jimmunol.179.2.984.

[83] Bruns HA, Schindler U, Kaplan MH. Expression of a Constitutively Active Stat6 in Vivo Alters Lymphocyte Homeostasis with Distinct Effects in T and B Cells. Journal of Immunology. 2003;170(7):3478–3487. doi:10.4049/jimmunol.170.7.3478.

[84] Kaplan MH, Sehra S, Chang HC, O’Malley JT, Mathur AN, Bruns HA. Constitutively Active STAT6 Predisposes toward a Lymphoproliferative Disorder. Blood. 2007;110(13):4367–4369. doi:10.1182/blood-2007-06-098244.

[85] Kuperman DA, Huang X, Koth LL, Chang GH, Dolganov GM, Zhu Z, et al. Direct Effects of Interleukin-13 on Epithelial Cells Cause Airway Hyperreactivity and Mucus Overproduction in Asthma. Nature Medicine. 2002;8(8):885–889. doi:10.1038/nm734.

[86] Grundberg E, Small KS, Hedman ÅK, Nica AC, Buil A, Keildson S, et al. Mapping Cis- and Trans-Regulatory Effects across Multiple Tissues in Twins. Nature Genetics. 2012;44(10):1084–1089. doi:10.1038/ng.2394.

[87] Westra HJ, Peters MJ, Esko T, Yaghootkar H, Schurmann C, Kettunen J, et al. Systematic Identification of Trans eQTLs as Putative Drivers of Known Disease Associations. Nature Genetics. 2013;45(10):1238–1243. doi:10.1038/ng.2756.

[88] Liang L, Morar N, Dixon AL, Lathrop GM, Abecasis GR, Moffatt MF, et al. A Cross-Platform Analysis of 14,177 Expression Quantitative Trait Loci Derived from Lymphoblastoid Cell Lines. Genome Research. 2013;23(4):716–726. doi:10.1101/gr.142521.112.

[89] Hao K, Bossé Y, Nickle DC, Paré PD, Postma DS, Laviolette M, et al. Lung eQTLs to Help Reveal the Molecular Underpinnings of Asthma. PLOS Genetics. 2012;8(11):e1003029. doi:10.1371/journal.pgen.1003029.

[90] Howell MD, Gao P, Kim BE, Lesley LJ, Streib JE, Taylor PA, et al. The Signal Transducer and Activator of Transcription 6 Gene (STAT6) Increases the Propensity of Patients with Atopic Dermatitis toward Disseminated Viral Skin Infections. The Journal of Allergy and Clinical Immunology. 2011;128(5):1006–1014. doi:10.1016/j.jaci.2011.06.003.

[91] Lee YL, Yen JJY, Hsu LC, Kuo NW, Su MW, Yang MF, et al. Association of STAT6 Genetic Variants with Childhood Atopic Dermatitis in Taiwanese Population. Journal of Dermatological Science. 2015;79(3):222–228. doi:10.1016/j.jdermsci.2015.05.006.

[92] Ferreira MAR, Matheson MC, Tang CS, Granell R, Ang W, Hui J, et al. Genome-Wide Association Analysis Identifies 11 Risk Variants Associated with the Asthma with Hay Fever Phenotype. The Journal of Allergy and Clinical Immunology. 2014;133(6):1564–1571. doi:10.1016/j.jaci.2013.10.030.

[93] Ferreira MAR, Jansen R, Willemsen G, Penninx B, Bain LM, Vicente CT, et al. Gene-Based Analysis of Regulatory Variants Identifies Four Putative Novel Asthma Risk Genes Related to Nucleotide Synthesis and Signaling. The Journal of allergy and clinical immunology. 2017;139(4):1148–1157. doi:10.1016/j.jaci.2016.07.017.

[94] Hedl M, Zheng S, Abraham C. The IL18RAP Region Disease Polymorphism Decreases IL-18RAP/IL-18R1/IL-1R1 Expression and Signaling through Innate Receptor–Initiated Pathways. The Journal of Immunology. 2014;192(12):5924–5932. doi:10.4049/jimmunol.1302727.

[95] Hirota T, Takahashi A, Kubo M, Tsunoda T, Tomita K, Sakashita M, et al. Genome-Wide Association Study Identifies Eight New Susceptibility Loci for Atopic Dermatitis in the Japanese Population. Nature genetics. 2012;44(11):1222–1226. doi:10.1038/ng.2438.

[96] Smyth DJ, Plagnol V, Walker NM, Cooper JD, Downes K, Yang JHM, et al. Shared and Distinct Genetic Variants in Type 1 Diabetes and Celiac Disease. The New England Journal of Medicine. 2008;359(26):2767–2777. doi:10.1056/NEJMoa0807917.

[97] Shrine N, Portelli MA, John C, Artigas MS, Bennett N, Hall R, et al. Moderate-to-Severe Asthma in Individuals of European Ancestry: A Genome-Wide Association Study. The Lancet Respiratory Medicine. 2019;7(1):20–34. doi:10.1016/S2213-2600(18)30389-8.

[98] Javierre BM, Burren OS, Wilder SP, Kreuzhuber R, Hill SM, Sewitz S, et al. Lineage-Specific Genome Architecture Links Enhancers and Non-Coding Disease Variants to Target Gene Promoters. Cell. 2016;167(5):1369–1384.e19. doi:10.1016/j.cell.2016.09.037.

[99] Zeller T, Wild P, Szymczak S, Rotival M, Schillert A, Castagne R, et al. Genetics and Beyond –The Transcriptome of Human Monocytes and Disease Susceptibility. PLOS ONE. 2010;5(5):e10693. doi:10.1371/journal.pone.0010693.

[100] Vibhuti A, Gupta K, Subramanian H, Guo Q, Ali H. Distinct and Shared Roles of *β*-Arrestin-1 and *β*-Arrestin-2 on the Regulation of C3a Receptor Signaling in Human Mast Cells. PLoS ONE. 2011;6(5). doi:10.1371/journal.pone.0019585.

[101] Zhang MJ, Xia F, Zou J. Fast and Covariate-Adaptive Method Amplifies Detection Power in Large-Scale Multiple Hypothesis Testing. Nature Communications. 2019;10(1):3433. doi:10.1038/s41467-019-11247-0.

[102] Lu Q, Powles RL, Wang Q, He BJ, Zhao H. Integrative Tissue-Specific Functional Annotations in the Human Genome Provide Novel Insights on Many Complex Traits and Improve Signal Prioritization in Genome Wide Association Studies. PLoS genetics. 2016;12(4):e1005947. doi:10.1371/journal.pgen.1005947.

[103] Lu Q, Powles RL, Abdallah S, Ou D, Wang Q, Hu Y, et al. Systematic Tissue-Specific Functional Annotation of the Human Genome Highlights Immune-Related DNA Elements for Late-Onset Alzheimer’s Disease. PLOS Genetics. 2017;13(7):e1006933. doi:10.1371/journal.pgen.1006933.

[104] Benjamini Y, Yekutieli D. The Control of the False Discovery Rate in Multiple Testing under Dependency. The Annals of Statistics. 2001;29(4):1165–1188. doi:10.1214/aos/1013699998.

[105] Thaventhiran JED, Lango Allen H, Burren OS, Rae W, Greene D, Staples E, et al. Whole-Genome Sequencing of a Sporadic Primary Immunodeficiency Cohort. Nature. 2020;583(7814):90–95. doi:10.1038/s41586-020-2265-1.

[106] Iotchkova V, Ritchie GRS, Geihs M, Morganella S, Min JL, Walter K, et al. GARFIELD Classifies Disease-Relevant Genomic Features through Integration of Functional Annotations with Association Signals. Nature genetics. 2019;51(2):343–353. doi:10.1038/s41588-018-0322-6.

