## Supplemental Figures for "Leveraging auxiliary data from arbitrary distributions to boost GWAS discovery with Flexible cFDR"

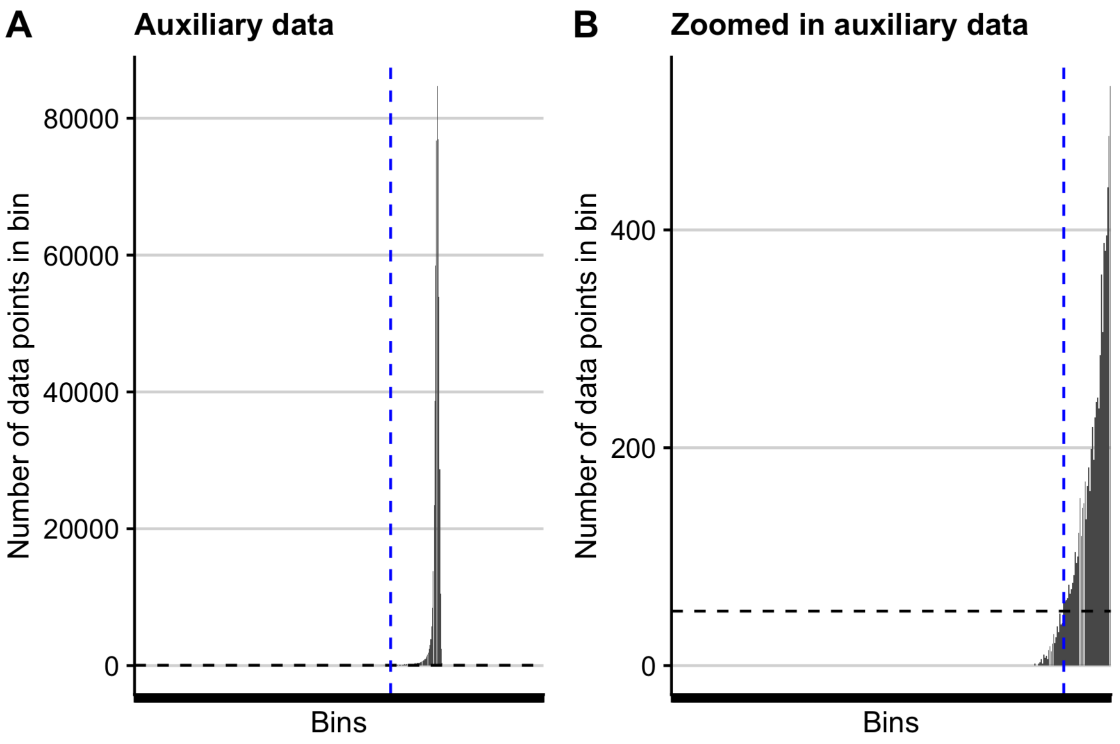


Figure S1: **Demonstration of left-censoring procedure.** Plots showing how many data points are in each grid space of the auxiliary data, $q$, over the support of the KDE for an example data set. (A) shows the full support of the KDE and (B) is zoomed in to the left tail. Black dashed line at $y = 50$ which is the default value of the `gridp` parameter in the `fcfdr::flexible_cfdr` function. Data points falling in grid spaces with fewer than 50 data points (those to the left of the blue dashed line) are left-censored, meaning that their value is replaced by the value of the left bound of the first grid space containing more than 50 data points. In practise, very few data points are left-censored.

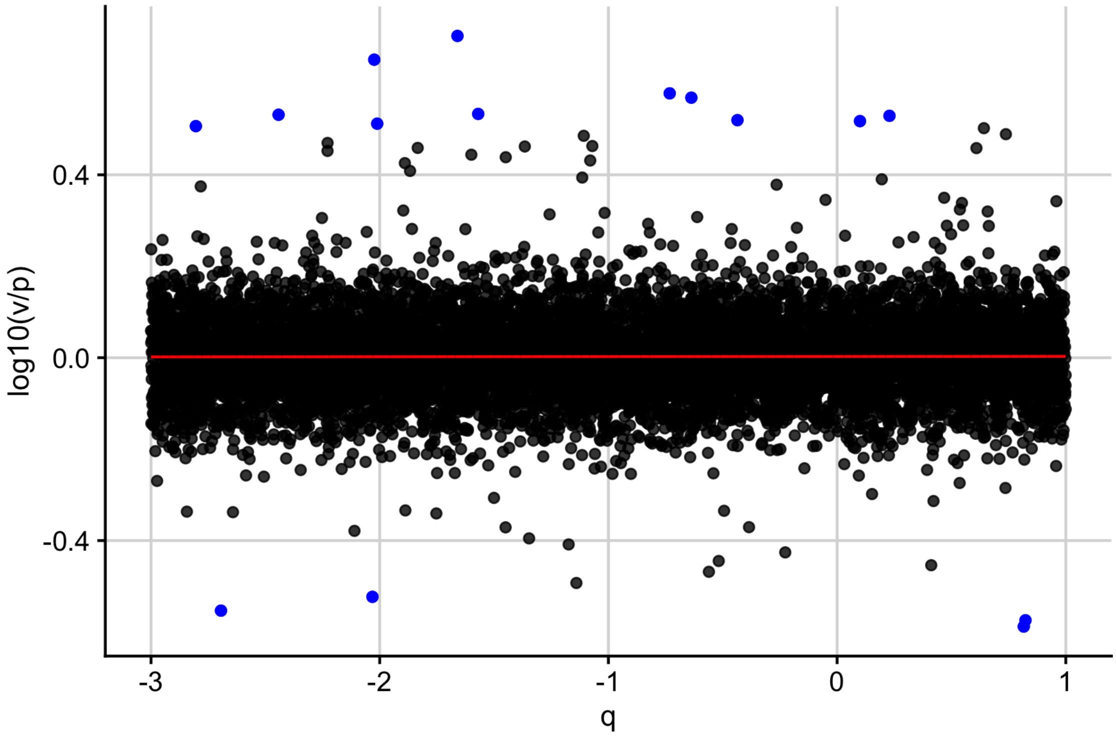


Figure S2: **Illustration of the spline correction procedure.** A spline with 5 knots is fitted to $log10(v/p)$ against $q$ using the 'bigsplines' R package (`https://cran.r-project.org/web/packages/bigsplines/index.html`). The distance between each data point and the fitted spline is calculated. If this distance is greater than the value of the 'dist_thr' parameter in the Flexible cFDR software (default value is 0.5), then the data point is mapped back to the spline and the corresponding $v$-value is recalculated using the fitted spline. In this example, the red line shows the fitted spline and the blue points are mapped back to the spline to generate new $v$-values.

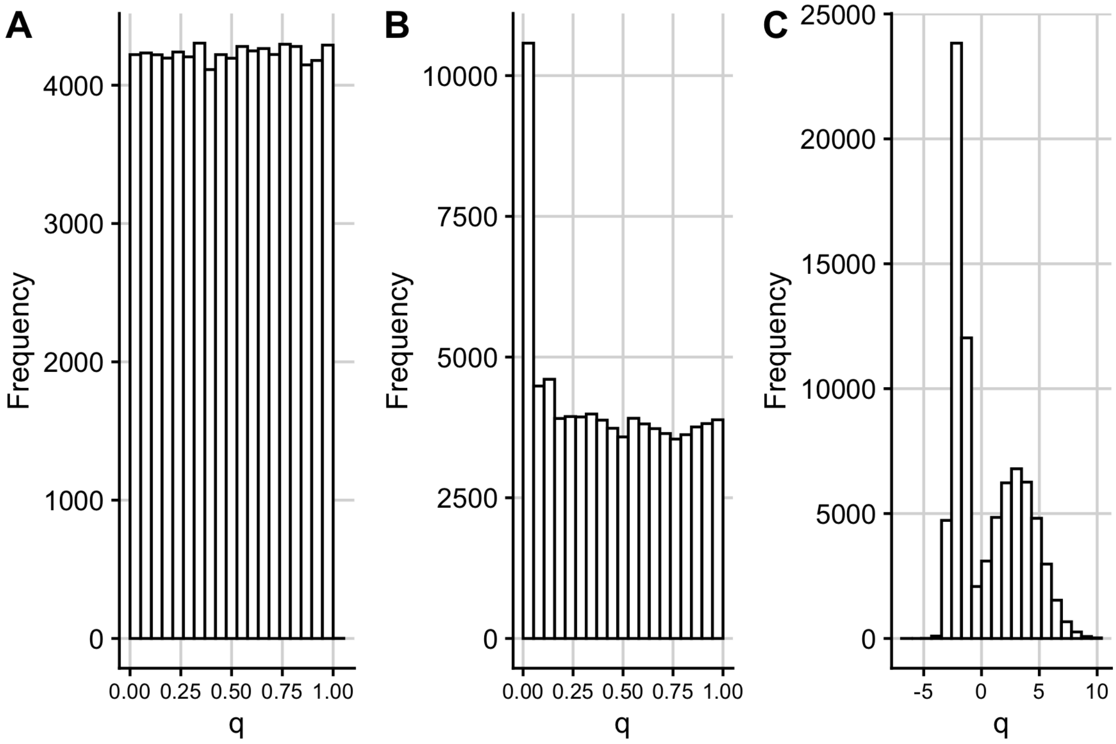


Figure S3: **Histograms of auxiliary data leveraged in simulation analysis.** (A) Example data leveraged in simulation A (simulated from standard uniform distribution). (B) Example data leveraged in simulation B (simulated $p$-values for related traits). (C) Example data leveraged in simulations C, D and E (simulated from a mixture normal distribution).

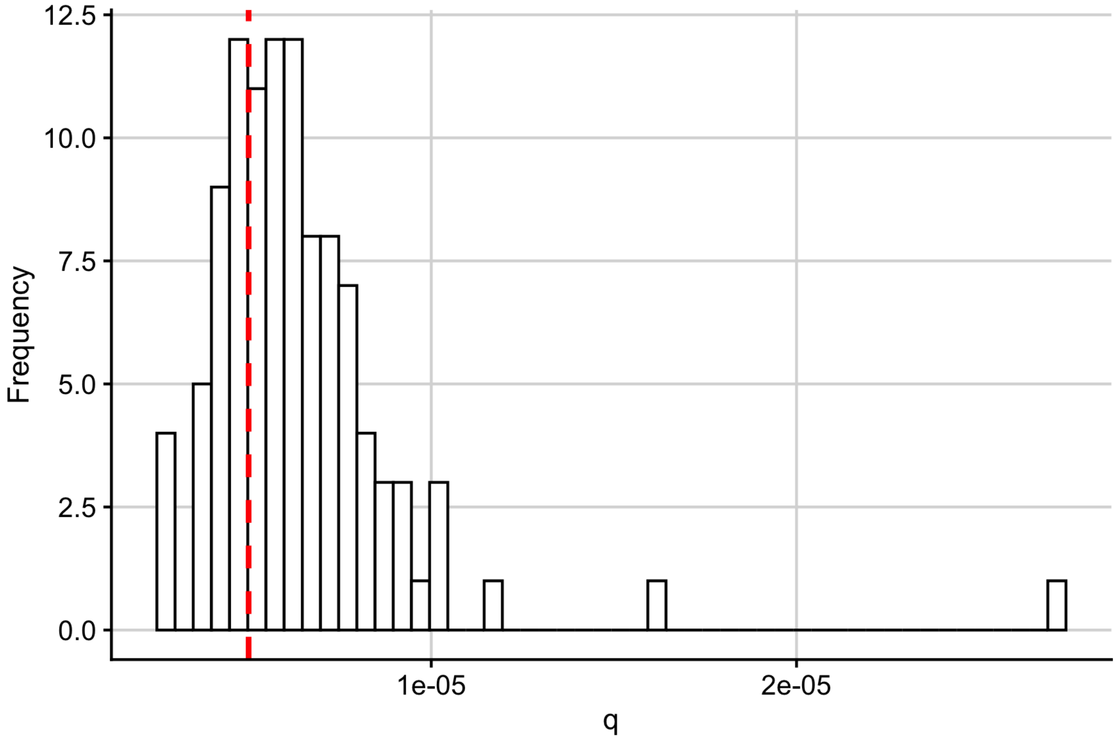


Figure S4: **Choosing an FDR threshold corresponding to the genome-wide significance p-value threshold in the simulation analysis.** Histogram of the maximum FDR-adjusted $p$-value (using BH method) amongst SNPs with $p \leq 5e-08$ in the simulation analysis. Red dashed line at the selected FDR threshold of $5e-06$.

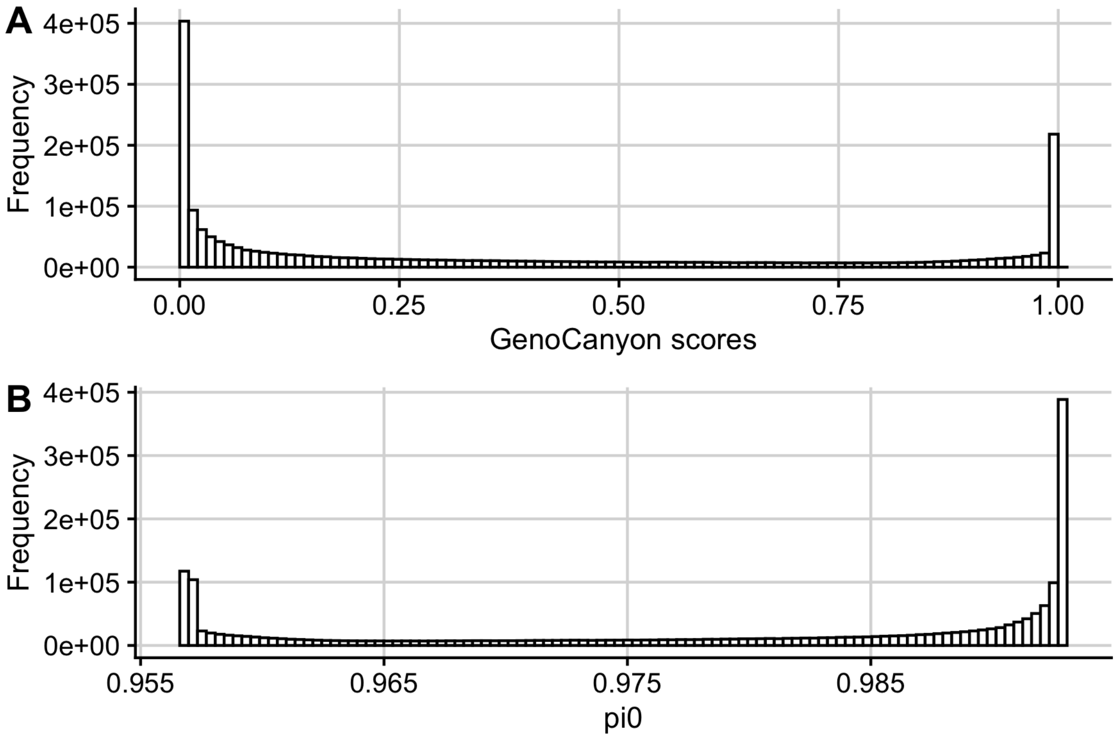


Figure S5: **Histograms from GenoCanyon application.** (A) Histogram of GenoCanyon scores for SNPs in the asthma GWAS data set. (B) Histogram of estimated pi0 values from BL.

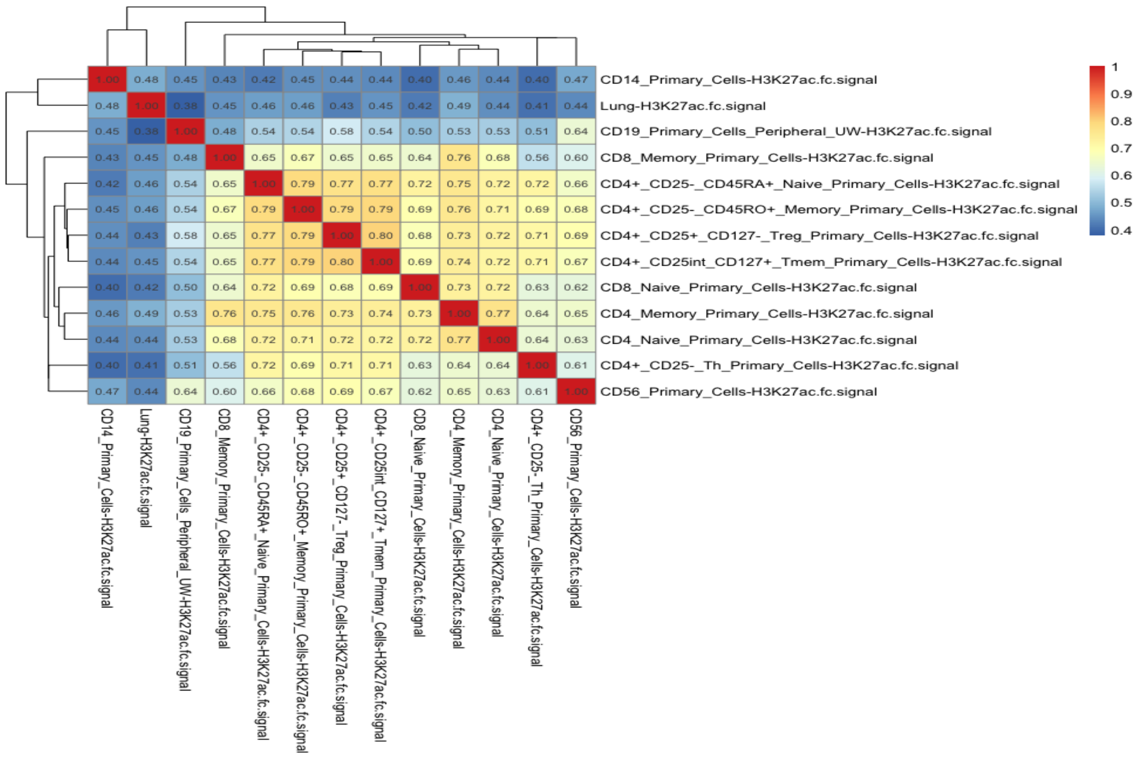


Figure S6: **Heatmap of the correlations between H3K27ac fold change values amongst asthma relevant cell types.**

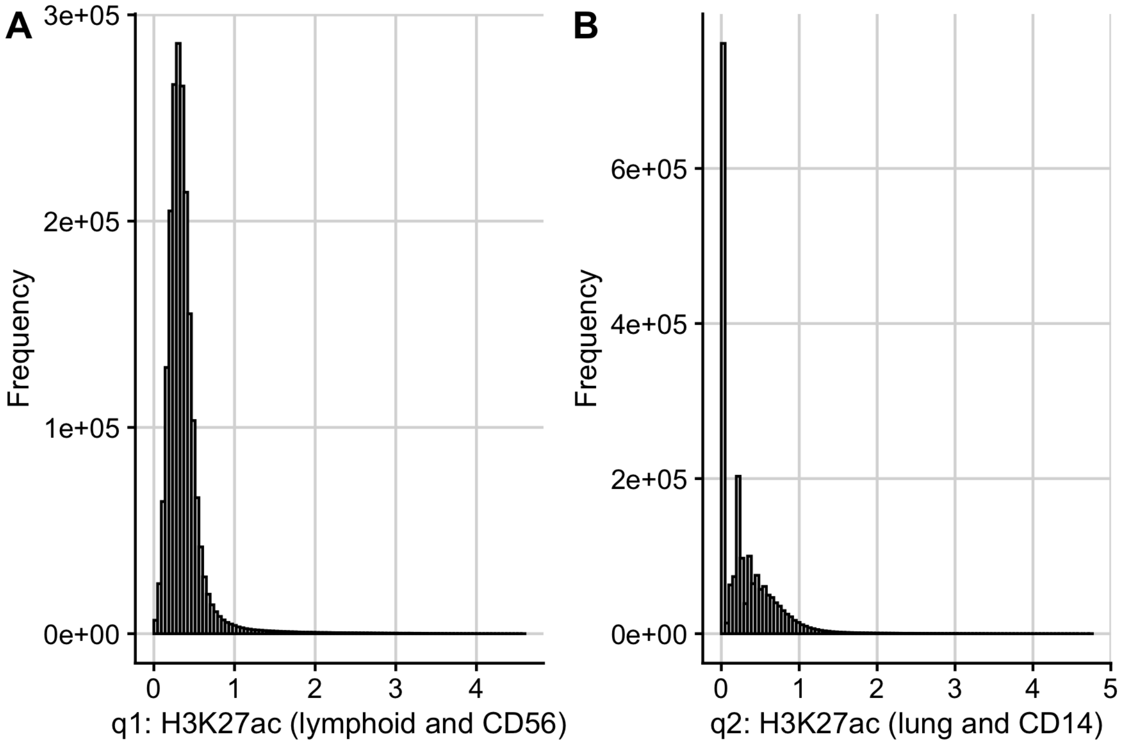


Figure S7: **Histograms of auxiliary data used in H3K27ac application.** (A) q1 is the average of (log transformed) H3K27ac fold change values in lymphoid and CD56 cell types (B) q2 is the average of (log transformed) H3K27ac fold change values in lung tissue and CD14+ cells.

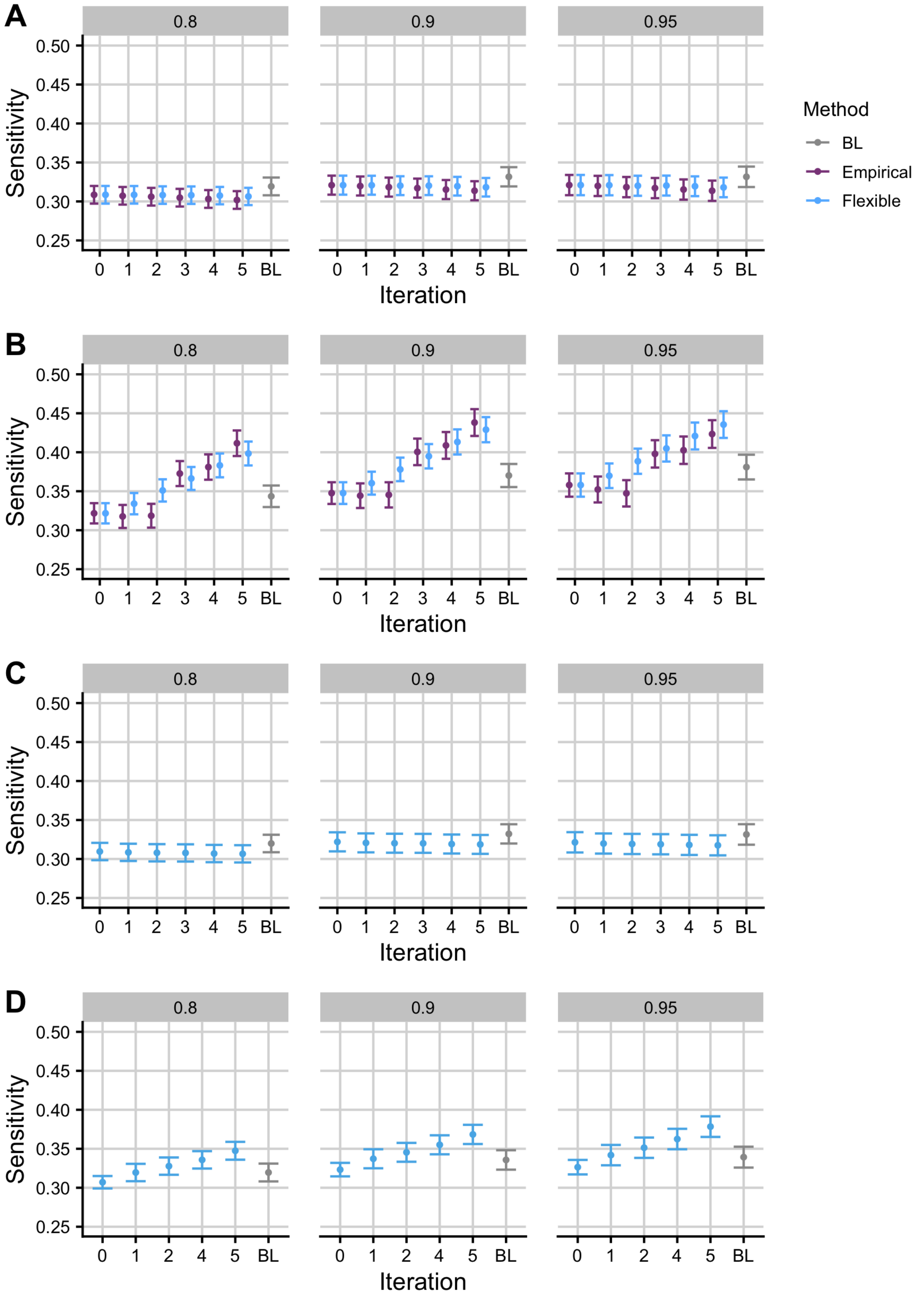


Figure S8: **Simulation results assessing the sensitivity when increasing the $r^2$ threshold used to call associated SNPs.** Mean +/- standard error for the sensitivity of FDR values from empirical and Flexible cFDR when iterating over independent (A; "simulation A") and dependent (B; "simulation B") auxiliary data that is bounded by $[0,1]$. Panels C and D show the results from Flexible cFDR when iterating over independent (C; "simulation C") and dependent (D; "simulation D") auxiliary data simulated from bimodal mixture normal distributions. BL refers to results when using Boca and Leek's FDR regression to leverage the 5-dimensional covariate data. Iteration 0 corresponds to the original FDR values. Our sensitivity proxy is calculated as the proportion of SNPs with $r^2 \geq X$ with a causal variant ("truly associated"), that were detected with a FDR value less than $5e-06$, where results are faceted for $X = 0.8, 0.9, 0.95$. Results were averaged across 100 simulations.

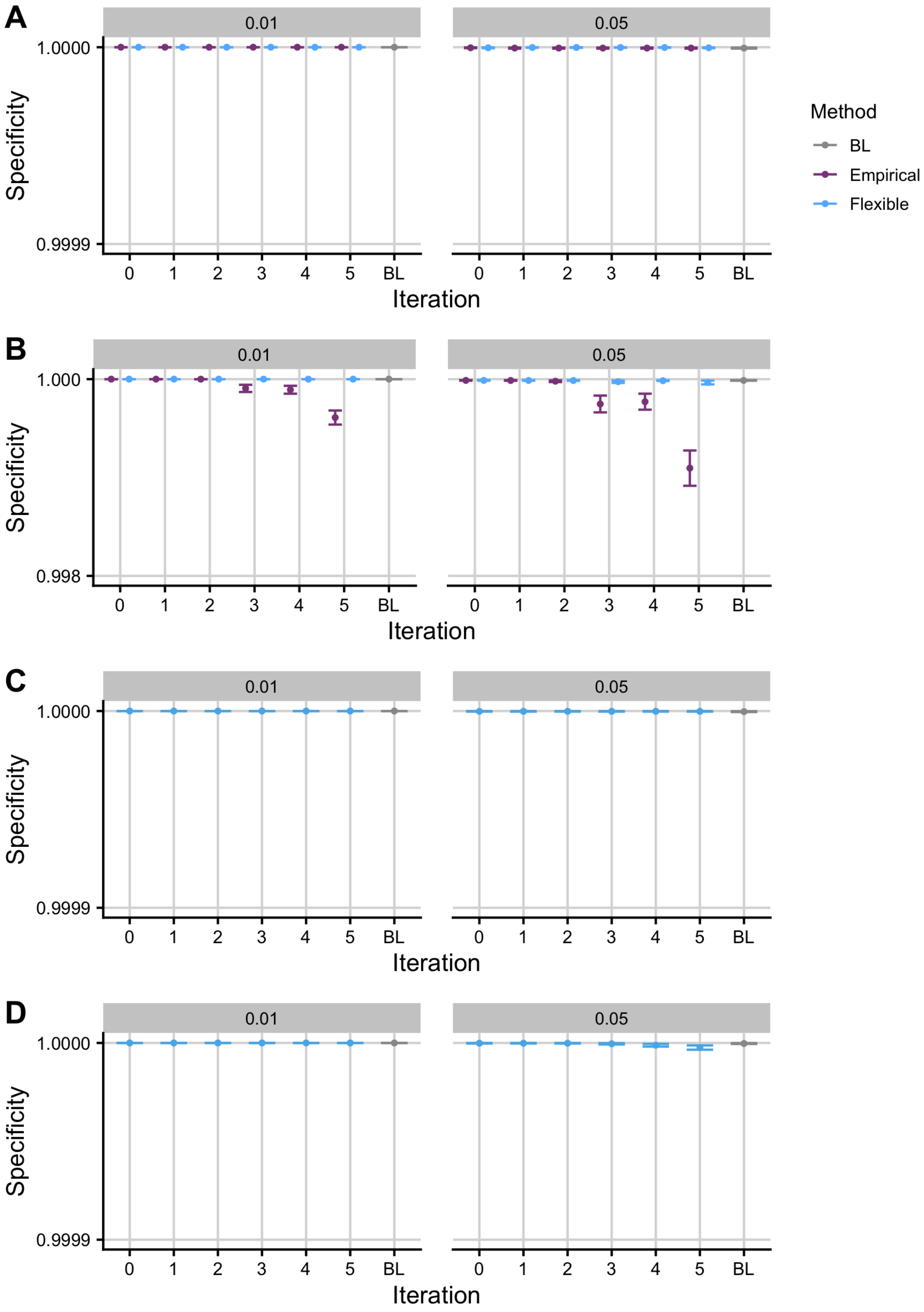


Figure S9: **Simulation results assessing the specificity when increasing the $r^2$ threshold used to call not-associated SNPs.** Mean +/- standard error for the specificity of FDR values from empirical and Flexible cFDR when iterating over independent (A; "simulation A") and dependent (B; "simulation B") auxiliary data that is bounded by $[0,1]$. Panels C and D show the results from Flexible cFDR when iterating over independent (C; "simulation C") and dependent (D; "simulation D") auxiliary data simulated from bimodal mixture normal distributions. BL refers to results when using Boca and Leek's FDR regression to leverage the 5-dimensional covariate data. Iteration 0 corresponds to the original FDR values. Our specificity proxy is calculated as the proportion of SNPs with $r^2 \leq X$ with all the causal variants ("truly not-associated"), that were not detected with a FDR value less than $5e-06$ , where results are faceted for $X = 0.01, 0.05$. Results were averaged across 100 simulations.

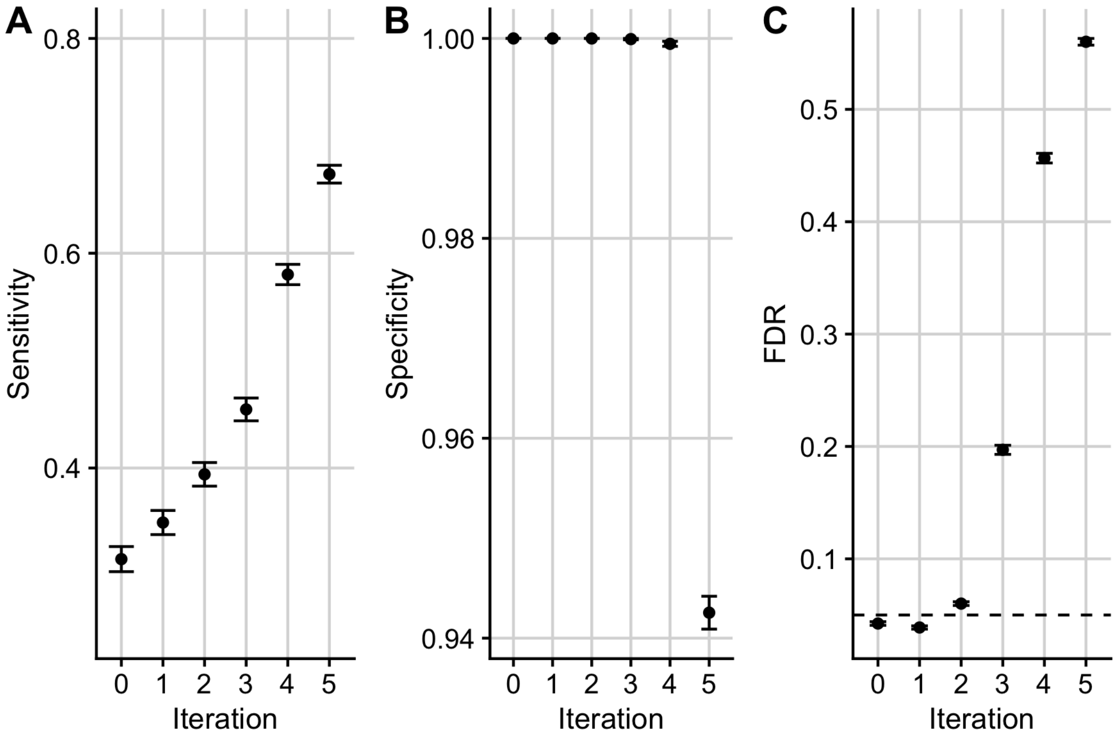


Figure S10: **Simulation results for using Flexible cFDR to iteratively leverage exactly the same auxiliary data.** Mean +/- standard error for the sensitivity (A) specificity (B) and FDR (C) of FDR values from Flexible cFDR when iterating over the same dependent auxiliary data ("simulation E"). Iteration 0 corresponds to the original FDR values. Our sensitivity proxy is calculated as the proportion of SNPs with $r^2 \geq 0.8$ with a causal variant (“truly associated”), that were detected with a FDR value less than $5e-06$. Our specificity proxy is calculated as the proportion of SNPs with $r^2 \leq 0.01$ with all the causal variants (“truly not-associated”), that were not detected with a FDR value less than $5e-06$. Our FDR proxy is calculated as the proportion of SNPs that were detected with a FDR value less than 0.05, that had $r^2 \leq 0.01$ with all the causal variants (“truly not-associated”) (we raised $\alpha$ to 0.05 in order to assess FDR control within a manageable number of simulations). Results were averaged across 1000 simulations.

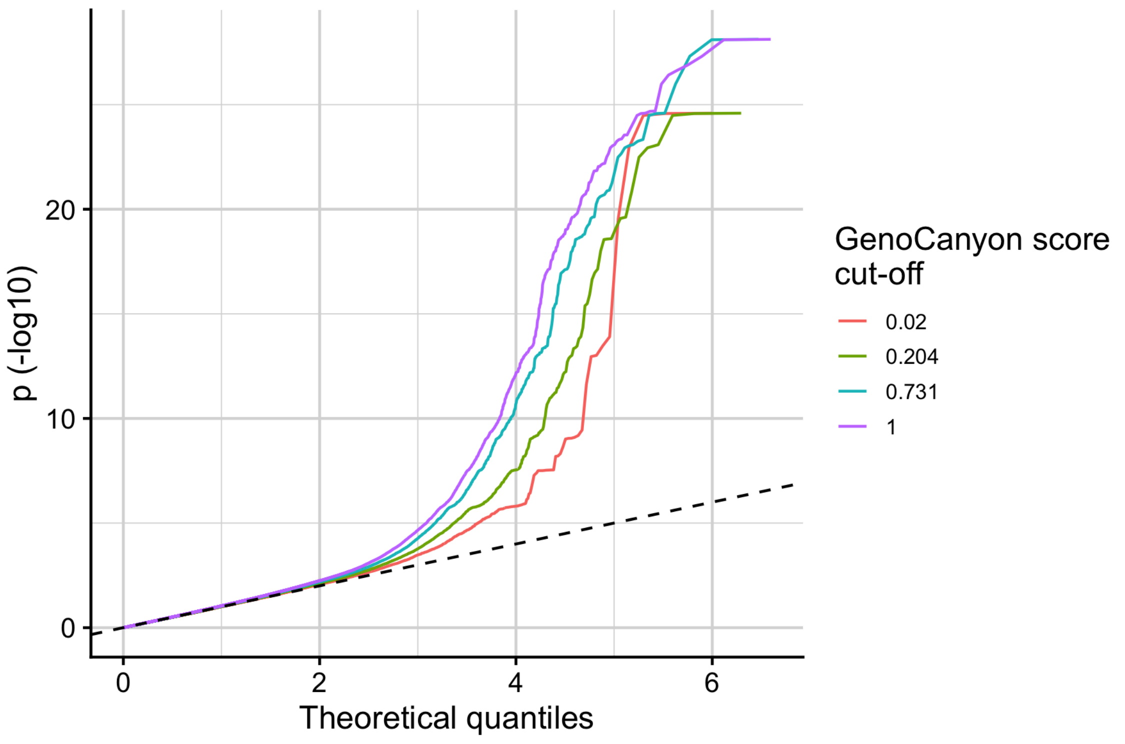


Figure S11: **Stratified Q-Q plot of empirical -log10 GWAS *p*-values for asthma against theoretical values stratified by GenoCanyon scores.** The values that were used to threshold the GenoCanyon scores are the quantiles of the distribution (0.020 is the 0.25 quantile, 0.204 is the 0.5 quantile, 0.731 is the 0.75 quantile and 1 is the maximum value).

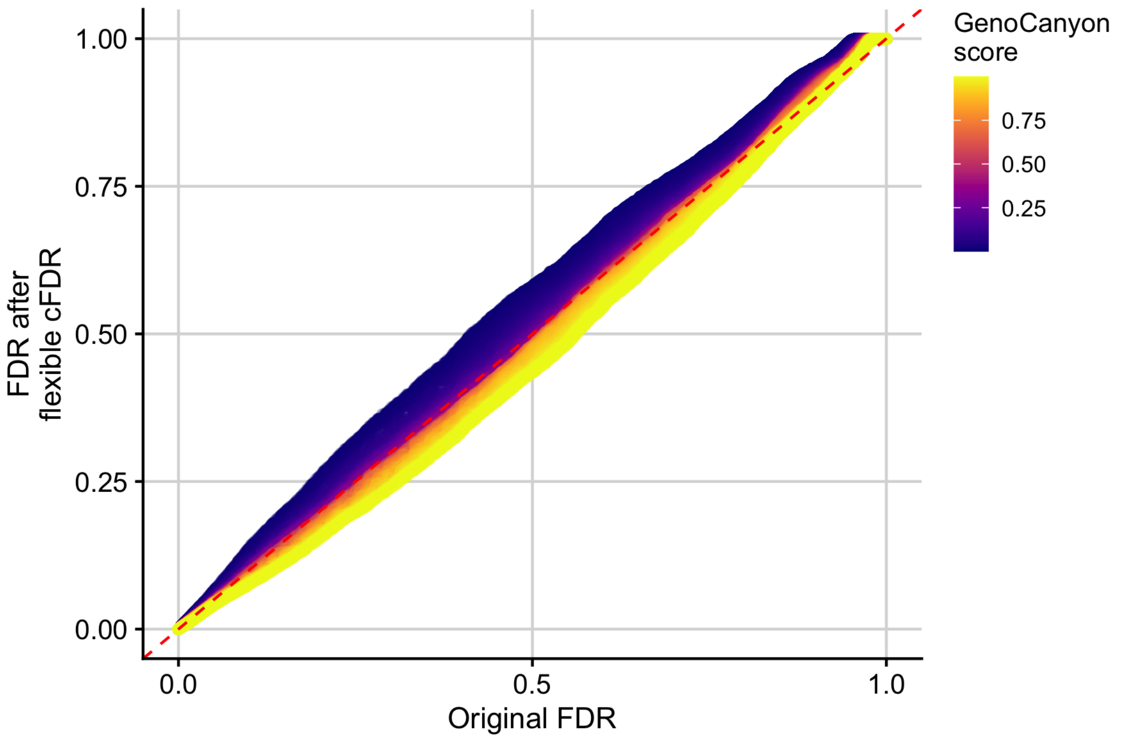


Figure S12: **Results when using Flexible cFDR to leverage GenoCanyon scores with asthma GWAS *p*-values.** FDR values after using Flexible cFDR to leverage GenoCanyon scores with asthma GWAS *p*-values against raw FDR values coloured by GenoCanyon score.

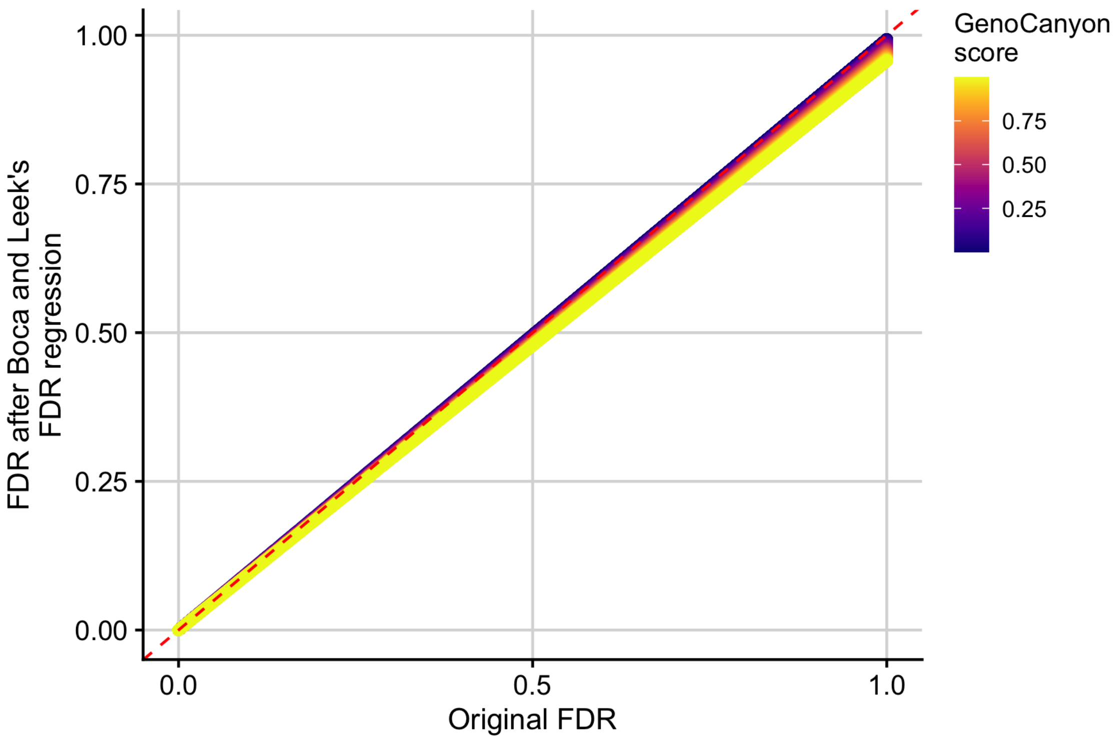


Figure S13: **Results when using BL to leverage GenoCanyon scores with asthma GWAS *p*-values.** Adjusted *p*-values from BL when leveraging GenoCanyon scores with asthma GWAS *p*-values against raw FDR values coloured by GenoCanyon score.

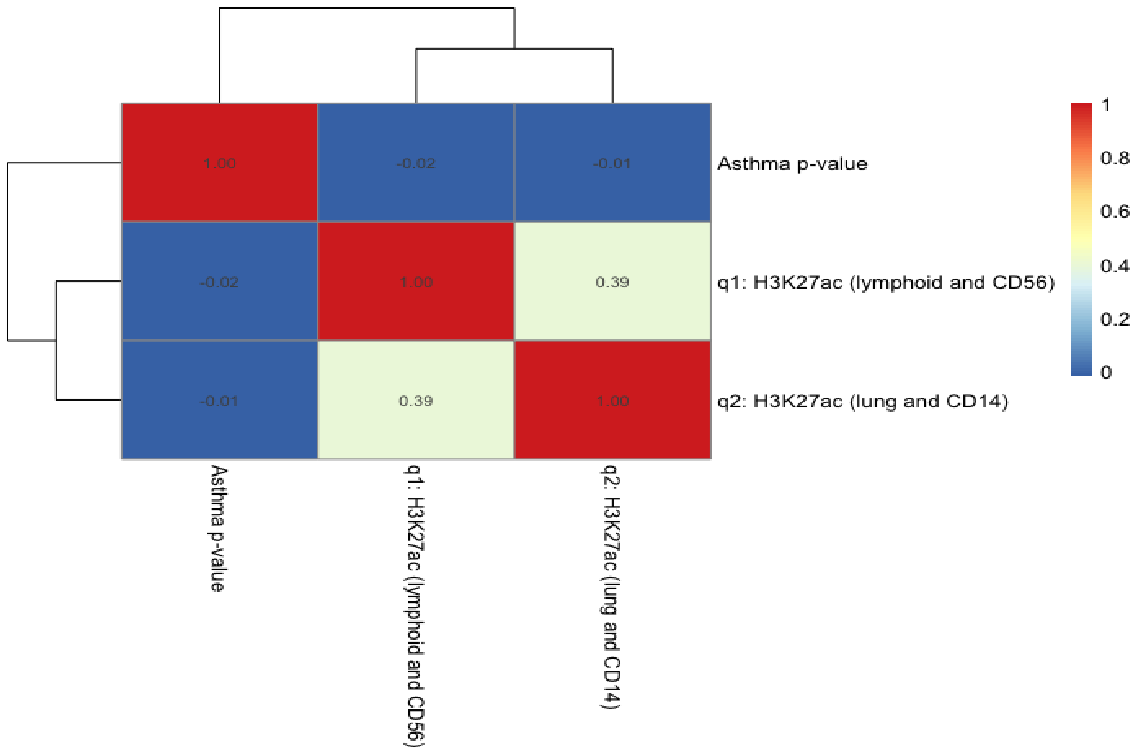


Figure S14: **Heatmap of correlations between asthma GWAS *p*-values and the summarised H3K27ac fold change values leveraged by Flexible cFDR.** q1 is the average of (log transformed) H3K27ac fold change values in lymphoid and CD56 cell types. q2 is the average of (log transformed) H3K27ac fold change values in lung tissue and CD14+ cells.

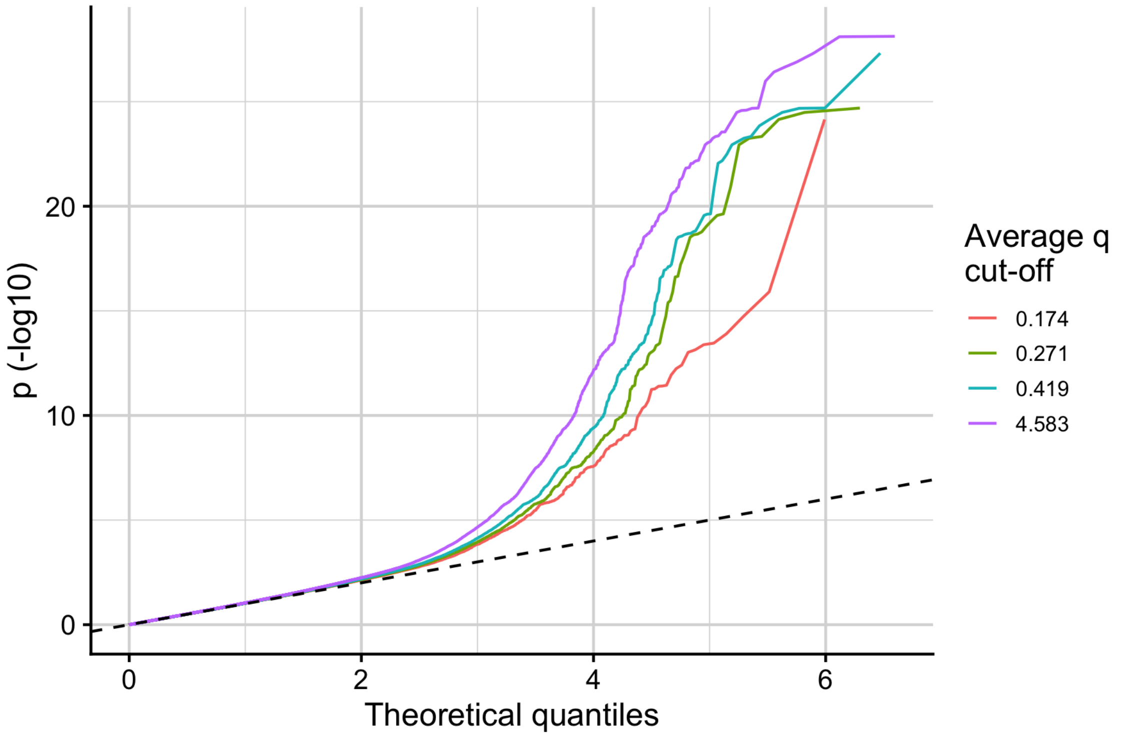


Figure S15: **Stratified Q-Q plot of empirical -log10 GWAS $p$-values for asthma against theoretical values stratified by average H3K27ac fold change values in asthma relevant cell types.** The values that were used to threshold $q$ (average H3K27ac fold change values) are the quantiles of the distribution (0.174 is the 0.25 quantile, 0.271 is the 0.5 quantile, 0.419 is the 0.75 quantile and 4.583 is the maximum value).

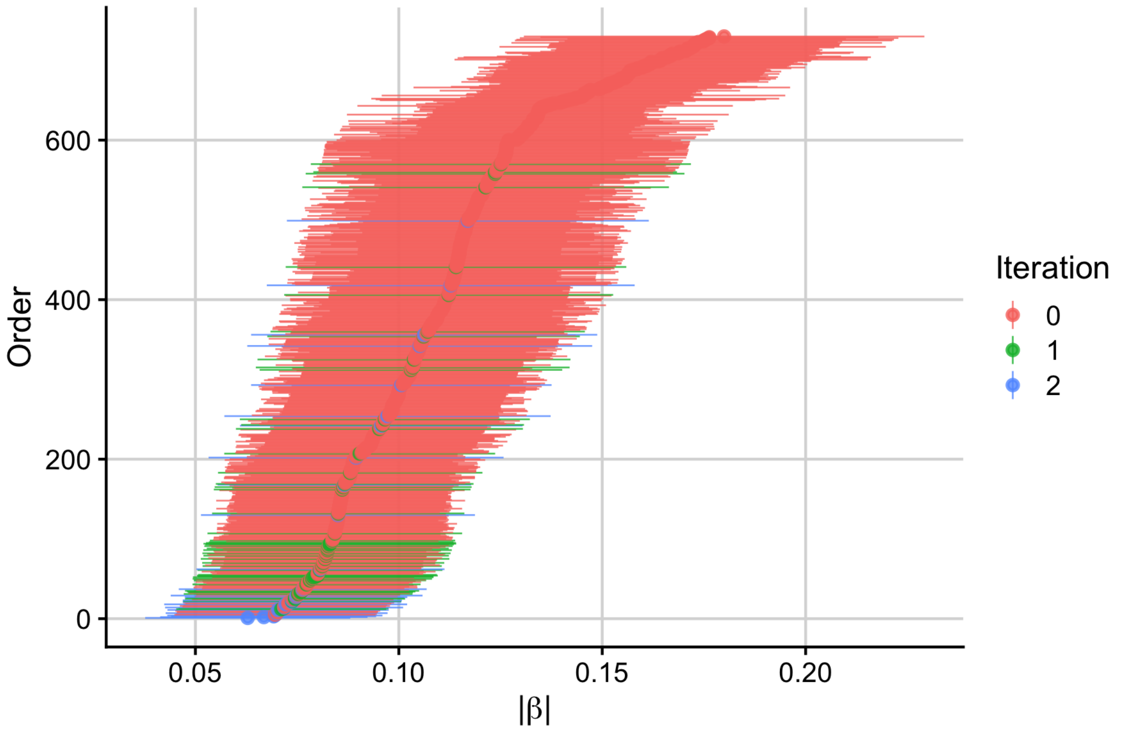


Figure S16: **Effect sizes of significant SNPs.** Absolute estimated effect sizes ($|\beta|$; log OR) $+/-1.96 \times SE$ of SNPs significantly associated ($FDR \leq 0.000148249$) with asthma in the original discovery GWAS data set ("iteration 0") and those newly significant after iteration 1 and 2 of cFDR.

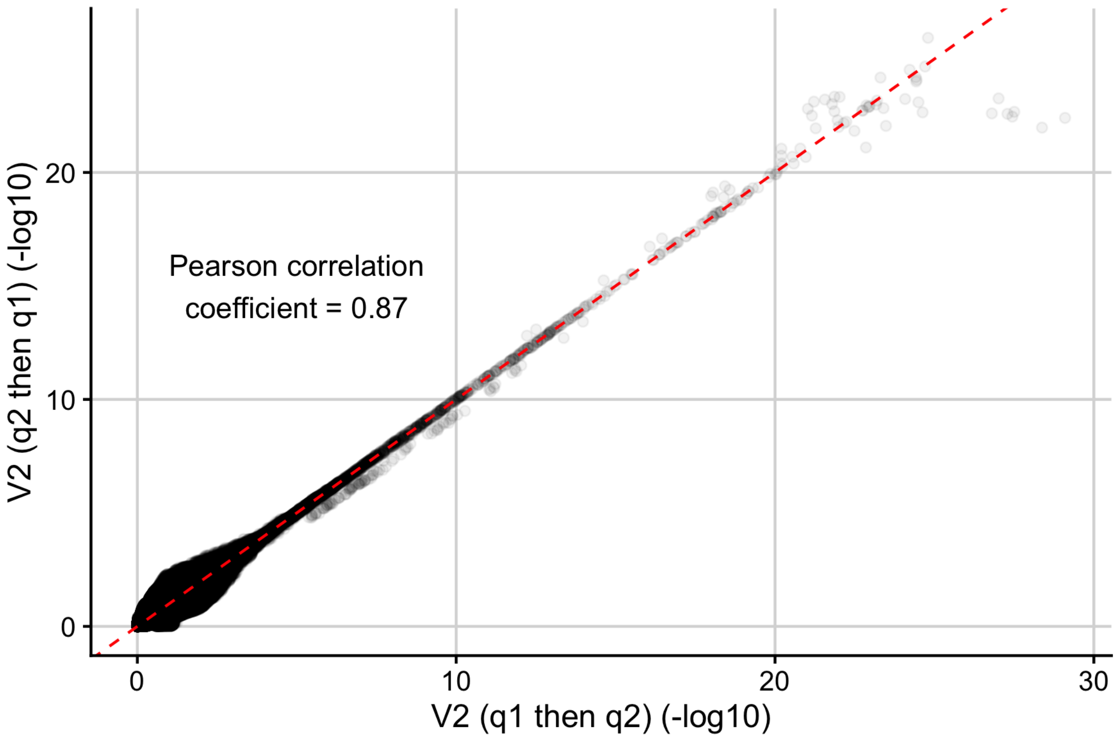


Figure S17: **Switching the order of iteration.** (-log10) $v$-values after 2 iterations of Flexible cFDR leveraging H3K27ac data when iterating over q2 and then q1 against (-log10) $v$-values when iterating over q1 then q2.

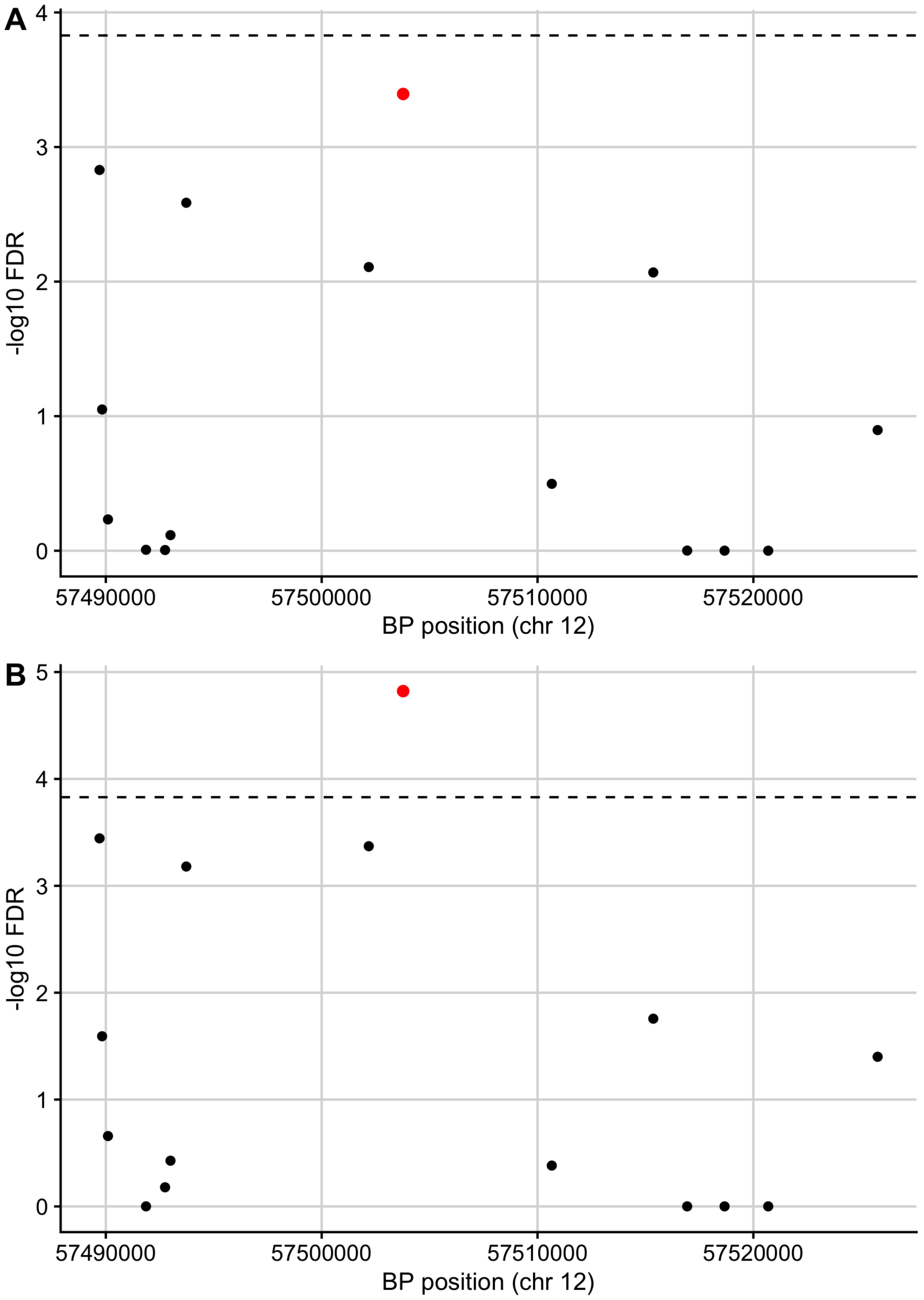


Figure S18: **Manhattan plots for genomic region containing the *STAT6* gene.** Manhattan plots of FDR values before (A) and after (B) applying Flexible cFDR for the region containing the *STAT6* gene (chr12:57489187-57525922). Black dashed line at FDR significant threshold. Red SNP is rs167769 (index SNP).

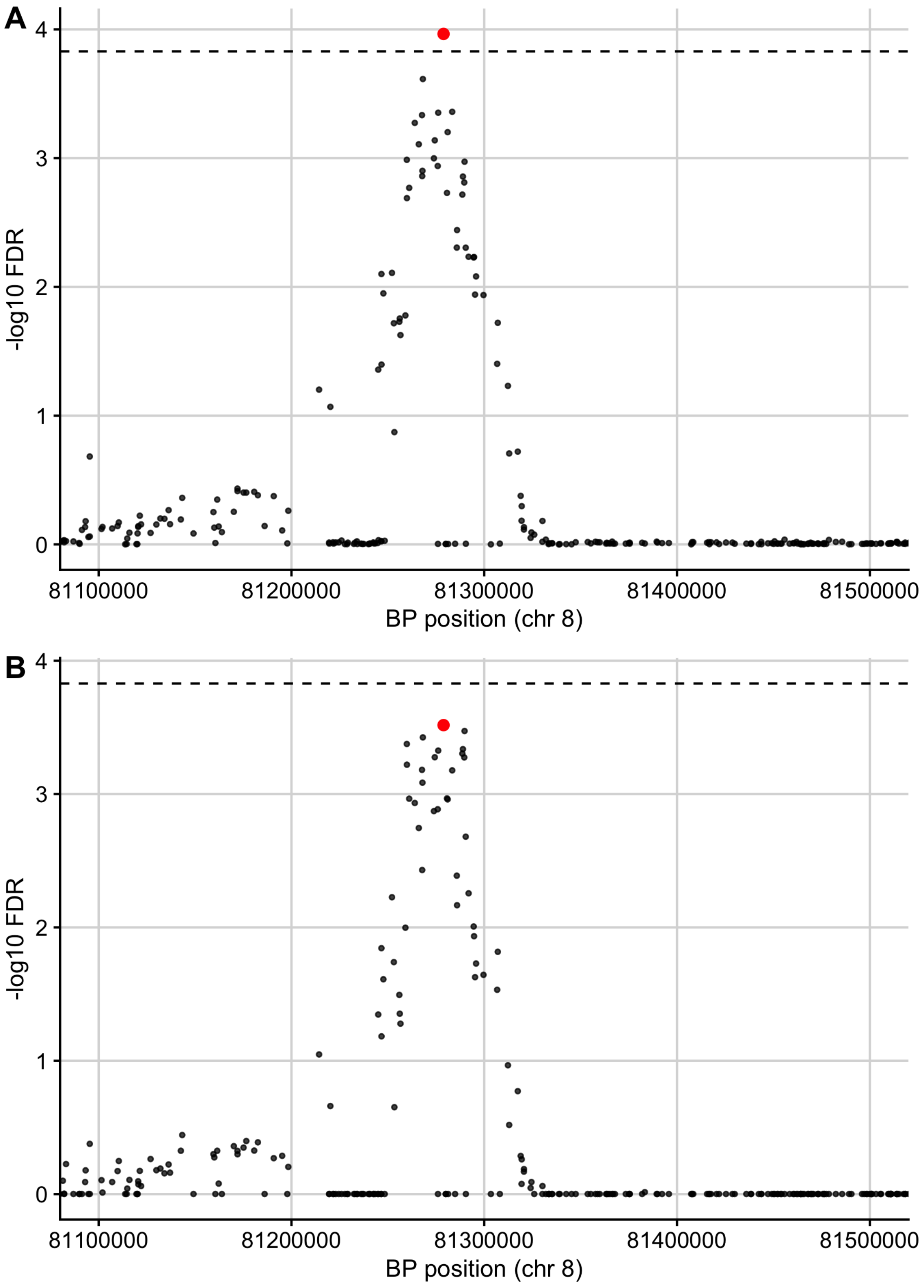


Figure S19: **Manhattan plots for region containing the index SNP rs12543811 that is no longer FDR significant after applying Flexible cFDR.** Manhattan plots of FDR values before (A) and after (B) applying Flexible cFDR for the region (chr8:81100000-81500000) containing index SNP rs12543811 (chr6:81278885) that is no longer FDR significant when applying Flexible cFDR.

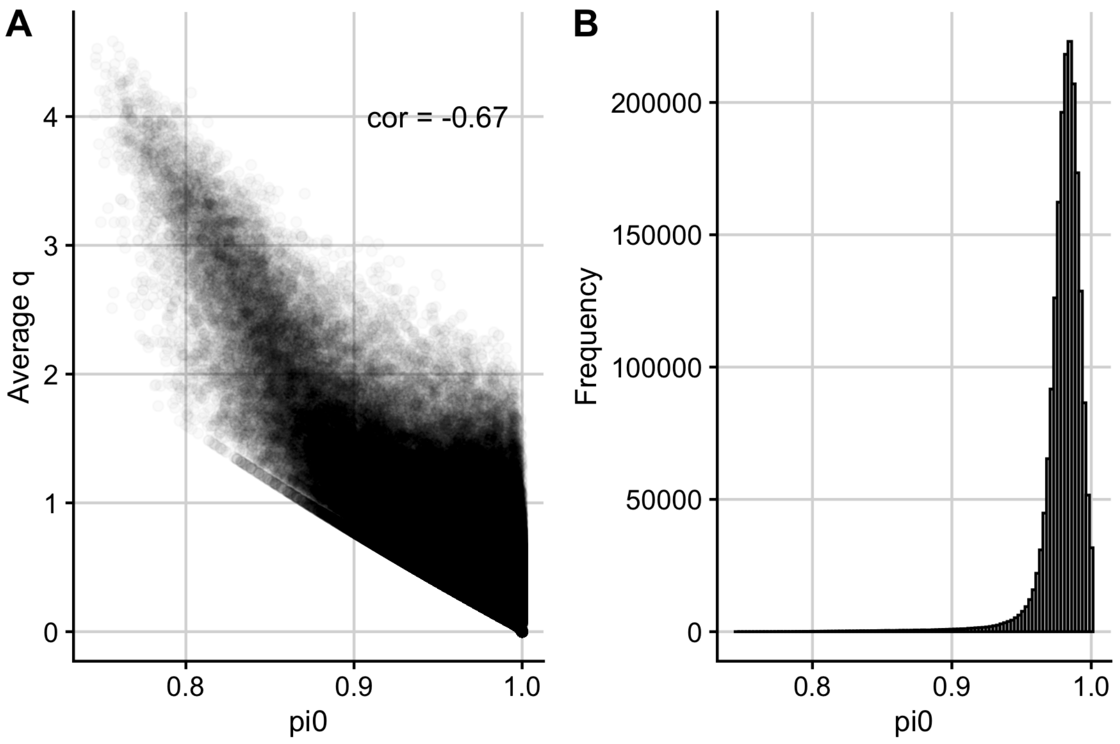


Figure S20: **Estimated pi0 values in BL when leveraging H3K27ac fold change values in relevant cell types with asthma GWAS *p*-values.** (A) Average H3K27ac fold change values in asthma relevant cell types ($q$) against estimated probabilities that the null hypothesis is true ('pi0') (B) Histogram of pi0 values for all 1,968,651 SNPs.
